# How to obtain cell volume from dynamic pH, temperature and pressure in plants

**DOI:** 10.1101/431700

**Authors:** Mariusz A. Pietruszka

## Abstract

We examined the pH/*T* duality of acidic pH and temperature (*T*) for the growth of grass shoots in order to determine the phenomenological equation of wall properties (‘equation of state’, EoS) for living plants. By considering non-meristematic growth as a dynamic series of ‘state transitions’ (STs) in the extending primary wall, we identified the ‘critical exponents’ (read: optimum) for this phenomenon, which exhibit a singular behaviour at a critical temperature, critical pH and critical chemical potential (μ) in the form of four power laws: *F_π_*(*τ*)∝|*τ*|*^β^*^−1^, *F_τ_*(*τ*)∝|*π*|^1−α^, *G_μ_*(*τ*)∝|*τ*|^−2−^*^α^*^+2^*^β^* and *G_τ_*(*μ*)∝|*μ*|^2−^*^α^*. The power-law exponents α and β are numbers that are independent of pH (or μ) and T, which are known as critical exponents, while π and τ represent a reduced pH and reduced temperature, respectively. Various scaling predictions were obtained – the convexity relation α + β ≥ 2 for practical pH-based analysis and a β ≡ 2 identity in a ‘microscopic’ representation. In the presented scenario, the magnitude that is decisive is the chemical potential of the H^+^ ions (protons), which force subsequent STs and growth. Furthermore, we observed that the growth rate is generally proportional to the product of the Euler beta functions of *T* and pH, which are used to determine the hidden content of the Lockhart constant Ф. It turned out that the evolution equation, when expressed in terms of the same dynamic set of variables, explains either the monotonic growth or periodic extension that is usually observed – like the one detected in pollen tubes – in a unified account. We suggest that cell growth evolves along the path with the least activity, thereby optimising growth under any physiological conditions. The pH dynamics in close-to-natural conditions appears to essentially be responsible for this extreme trajectory, thereby providing a highly nonlinear pH(*t*), 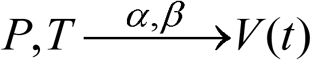 transformation. Moreover, the drops in pH that are induced by auxin or fusicoccin, when next converted by the augmented Lockhart equation, are enough to explain a significant fraction of the increase in the growth rate. A self-consistent recurring model is proposed to embrace the inherent complexity of such a biological system, in which several intricate pathways work simultaneously, in order to reconcile the conflicting views of plant cell extension and growth. Eventually, we pose the question: Is the chemical potential of protons a master regulator for tip-growing cells?

**Author summary:** In plant development, sudden changes such as cell expansion or pollen tube oscillations seem to depend on a correlative group of events rather than on slow shifts in the apex. Hence, in order to understand or to control the processes in the extending cell wall, we need to unravel the general principles and constraints that govern growth. The quest for these principles has primarily focused on the molecular, though merely descriptive, level. Here, we show that it is possible to analyse oscillatory state changes computationally without even requiring knowledge about the exact type of transition. Our results suggest that the cell wall properties and growth of plant cells can be accurately and efficiently predicted by a set of physical and chemical variables such as temperature, pressure and the dynamic pH of the growing plant, which build a scaffold for more specific biochemical predictions. In this context, we observed that cell growth evolves along the path the least action, thereby optimising growth under any physiological conditions. The model equations that we propose span the fields of the biological, physical, chemical and Earth sciences. The common denominator that ties the growth factors together is the chemical potential of protons, which is possibly a central core-controlling mechanism that is able to produce a macroscopic outcome, i.e. structurally and temporally organised apical growth.

## PART I Equation of Wall Properties for Plants

### Introduction

#### General outline

This part deals with the derivation and determination of an ‘equation of state’ (EoS) or – equivalently – an equation of wall properties for plants that integrates the relationship between temperature and pH (or chemical potential, μ) with growth. Based on reliable (published) data and our own experimental data, we sought to identify the ‘state’ variables that are necessary to describe optimal growth. The limited number of parameters can be misleading; this approach does not oversimplify the complexity of biological systems, but conversely, takes all of them into account by building an ‘outer scaffold’ that cannot be surpassed. We cannot dismiss the possibility that this is the role of phenomenology in making universal statements, in this case imposing a physical constraint (i.e. EoS) on the extending primary wall.

A commonly held view is that biological systems are complex (and molecular biologists are deluged with data), and that therefore modelling has to cope with the fact that the degrees of freedom are numerous, and correspondingly, the number of parameters should be high. Though physical systems on a microscopic scale are also complex and the number of degrees of freedom in real systems is abundant (e.g. Avogadro number, as in the solid state physics), their behaviour can usually be described by a few parameters (Ginzburg, 2004) within the framework of a phenomenological theory. A similar argument also concerns quantum models (Hubbard, 1963), which have only two parameters: *t* – for the “hopping” integral and *U* – for the Coulomb interaction of electrons within narrow energy bands. In what follows, I argue that, similar to physical systems, a low number of relevant parameters may also be sufficient to describe complex biological systems. To support this view let me quote Portes et al. (2015): “This is not to invalidate the strive for detailed models, which are necessary to understand the role of specific components of interest, however, they require a compatible amount of data. In contrast, more general models help to unveil the fundamental principles driving a phenomenon of interest, helping to identify the key regulatory interactions, albeit lacking details”. In what follows, we have chosen the second route.

The growth of plant cells and organs (like extending cylindrical organs such as grass shoots) are affected by light and humidity, which along with pH and temperature are the most important factors that influence growth. It can be a rewarding enterprise to understand plant growth in terms of the time evolution of STs (e.g. from an ordered to a disordered state, from a stressed to a relaxed state, from a “covalent bonds state” to a “disruption of covalent bonds state”, etc.) that take place in the peripheral cell wall. Such a transition (‘state transition’, ST, or equivalently ‘state change’, which should <u>not</u> be confused with the typical *phase transition* that is encountered in physics, since we are not at an equilibrium steady state) occur *via* the exchange of particles and energy-consuming metabolic processes (such as energy-conserving ATP) with the inside of cell compartment – the vacuole and cytoplasm, which are treated as the reservoirs of molecules and heat (thermostat). Hence, we consider living organisms to be *open systems* that dissipate energy and exchange entropy (heat) and matter (molecules) with the environment (Barbacci et al., 2015), a thermostat. This process is accomplished by the presence of pH- and *T*-dependent Euler beta functions in model equations, which represent the modes of interaction (like heat and mass transfer) with the outside world of a growing plant. Even though we considered a single ST, a cascade (ratchet) of subsequent chemical potential – induced STs, which may constitute the extension of the cell wall and growth, can be imagined (Part II). We demonstrate the prevailing role of the chemical potential and propose that the growth of plant cells and organs is regulated, at the molecular level, by the chemical potential (Matlak et al., 2004; van der Marel 2004) of H^+^ ions. We also show, at a phenomenological level, that the ionic fluxes may generate power-law relations. The proposed model correctly accounted for the set of experimental results and predicted the growth of grass shoots from pH with a high fidelity.

#### Preliminaries

Plant developmental systems have evolved within the universal limitations that are imposed by the plant cell wall (Lintilhac, 2014). Modulation of the mechanical properties occurs through the control of the biochemical composition and the degree and nature of interlinking between cell wall polysaccharides (Bidhendi and Geitmann, 2015). Plant cells encase themselves in a complex polysaccharide wall (Cosgrove, 2005a) and characteristically obtain most of their energy from sunlight *via* the photosynthesis of the primary chloroplasts. The expansive growth of turgid cells, which is defined as an irreversible increase in cell volume, can be regarded as a physical process that is governed by the mechanical properties of the cell wall and the osmotic properties of the protoplast (Schopfer, 2006). The precise biochemical mechanism that regulates the ability of the growth-limiting walls to extend irreversibly under the force of turgor pressure has not yet been identified (Kutschera, 2000), but it is correlated with the loosening of cells walls, which increases the cell’s susceptibility to expansion.

Growing plant cells characteristically exhibit acid growth (Rayle and Cleland, 1970; Hager et al., 1971; Cosgrove, 1989), which has been formulated in the form of the “acid growth hypothesis” (Hager, 2003; Taiz and Zeiger, 2006). The acid growth hypothesis was proposed over 40 years ago and is widely considered to play a role in expansive cell growth. The acid growth hypothesis postulates that both phytotoxin fusicoccin (FC) and the growth-promoting factor auxin (indole-3-acetic acid, IAA) cause wall-loosening and produce the concomitant induction of growth (growth enhancement) through the rapid acidification of the extension-limiting cell wall (Cleland, 1973; Kutschera, 1994). These enigmatic “wall-loosening processes” are, in fact, minor changes within the polymer network of the extension-limiting walls (i.e. the incorporation of proteins, the enzymatic splitting of polymer backbones or covalent cross-links or the disruption of non-covalent interactions between wall polymers *via* expansin activity (McQueen-Mason and Cosgrove, 1994). Molecular studies revealed that expansins are a large family of proteins with 38 members in *Arabidopsis thaliana*, which can be divided into three subfamilies: α-, β- and γ-expansins (Li et al. 2002). It was shown that individual expansins might be tissue- or process-specific, and that a loss-of-function mutation resulted in a disorder in a specific aspect of plant growth or development, for example in root hair development (reviewed by Marzec et al., 2015a).

Growth is accomplished through the enlargement of the cell volume owing to water uptake, which maintains the appropriate inside pressure in the vacuole as well as the irreversible extension of the pre-existing (primary) cell wall. Hereafter, we will assume an almost constant or slowly varying turgor pressure, which by definition is a force that is generated by water pushing outward on the plasma membrane and plant cell wall that results in rigidity in a plant cell and the plant as a whole.

Expansive growth is the result of the coupling effects (Barbacci et al., 2013) between mechanical (pressure), thermal (temperature) and chemical energy (pH). The thermal sensitivity of biochemical processes refers to the coupling effect between the thermal and chemical energies. In the search for a plant-wall specific EoS in growing biological cells or non-meristematic tissues, we considered two ‘state variables’, namely physiological temperature T (from 0 to 45 degrees in the Celsius scale) and pH at a constant turgor. We note that pH is not *sensu stricto* a fundamental physical quantity, and therefore, cannot be treated as a usual intensive variable. Nonetheless, relying on the definition of pH and then considering the chemical potential (Baierlein, 2001), we can propose the following.

Evidence has accumulated that the final goal of auxin action (Steinacher et al., 2012; Lüthen, 2015 for review; Majda and Robert, 2018) is to activate the plasma membrane (PM) H^+^-ATPase, a process that excretes H^+^ ions into the cell wall. The auxin-enhanced H^+^-pumping lowers the cell wall pH, activates pH-sensitive enzymes and proteins such as expansins (Cosgrove, 1993), xyloglucans (Fry et al., 1992) or yieldins (Okamoto-Nakazato et al., 2001) within the wall and initiates cell-wall loosening and extension growth. It has also been observed that when auxin-depleted segments were submerged in an acid buffer solution, the segments started their elongation growth immediately (“acid growth”), whereas the auxin-induced growth (“auxin growth”) began after a delay (lag phase), as is shown in Fig. 1 (curve 1) in Hager (2003); see also the remaining plots, which are of quite a different character, (curves 2–6) that correspond to “acid growth”. From the many investigations that have been done for more than four decades, it has been deduced that protons, which are extruded into the wall compartment, are directly responsible for the wall-loosening processes through the hydrolysis of covalent bonds, transglycosylation or the disruption of non-covalent bonds (thus changing the ‘state’ of the system). The growth effect is illustrated in Fig. 2 in Hager (2003), where the “Zuwachs” (increments in %) is plotted against pH.

**Figure 1.**
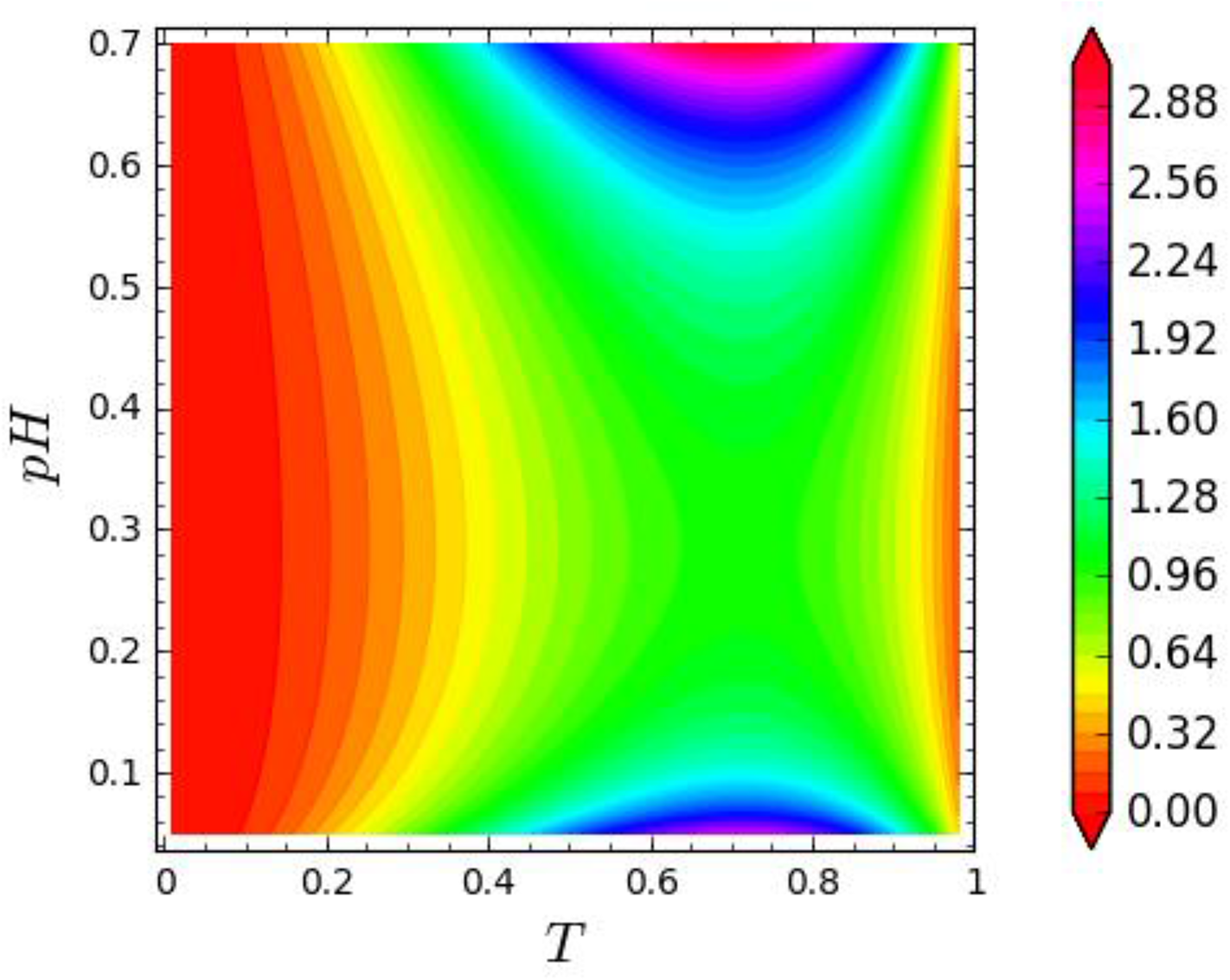
Plot of the ‘state function’ *F* given by equation (I-3) showing the optimum conditions for plant cell/organ growth. Both coordinates, temperature (*T*) and acidity or basicity (alkaline medium) pH, are rescaled to unity. The best conditions for maximum growth (green) are found at the saddle point (a stationary point). The simulation parameters that were used are similar to those that were used for auxin-induced growth – see SI Table 1.

**Figure 2.**
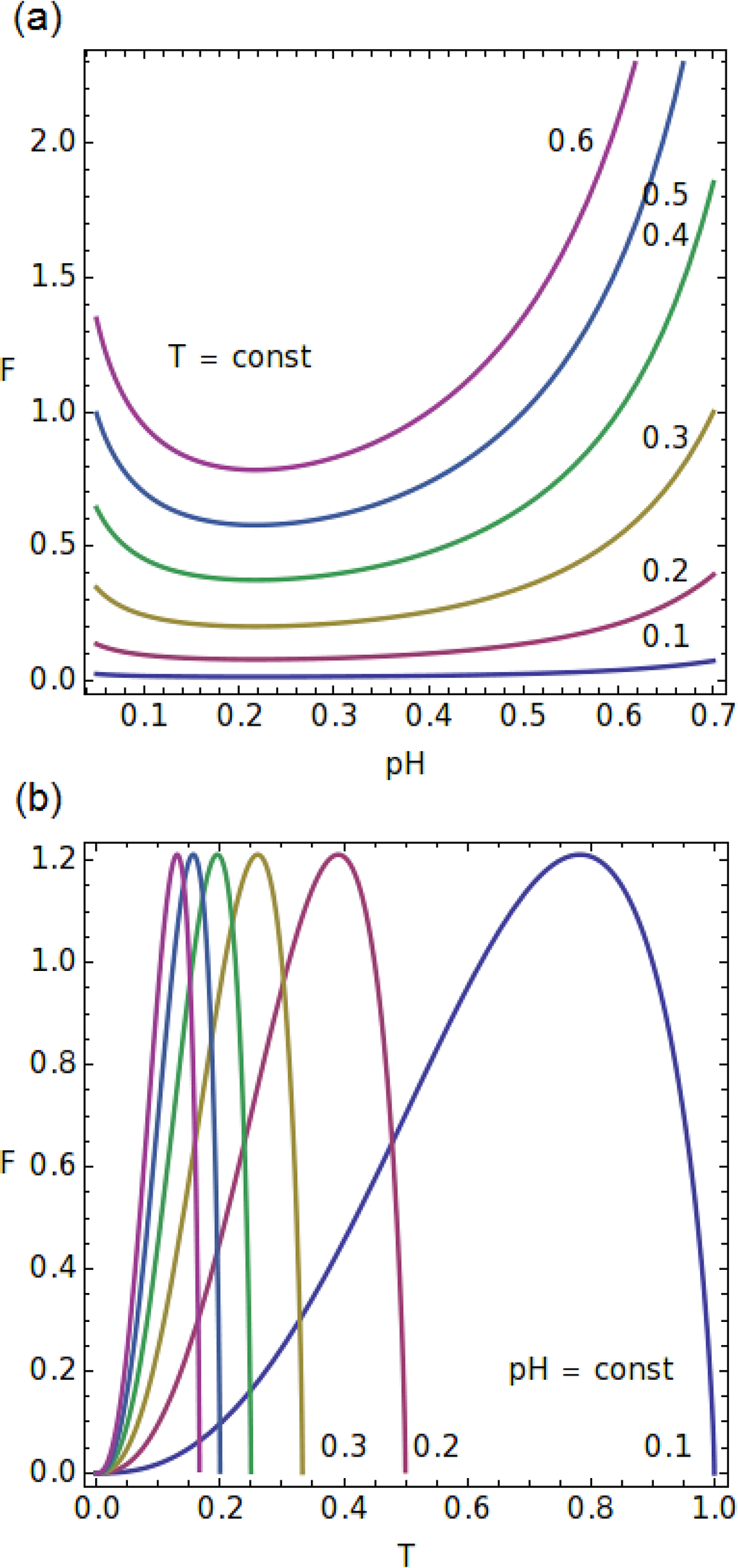
The (a) isotherms (scaled values indicated) as a function of pH (scaled) of the function *F*, Eq. (I-3); (b) constant pH curves as a function of temperature (scaled). Simulation parameters: α = 1.7 and β = 3.52 (SI Table 1).

However, pH can be expressed in terms of a more elementary quantity, namely the chemical potential μ_H+_ of protons. Note that indirect measurements of the chemical potential by means of pH can be compared to our previous measurements of the electromotive force (EMF) in order to detect phase transitions, which are localised by the ‘kinks’ in the chemical potential in condensed matter physics (Matlak and Pietruszka, 2000; Matlak et al. 2001). Although pH is not a typical generalised coordinate, the microscopic state of the system can be expressed through it in a collective way.

Temperature is among the most important environmental factors that determine plant growth and cell wall yielding (Pietruszka et al., 2007). Plants are responsive to temperature and some species can distinguish differences of 1°C. In Arabidopsis, a warmer temperature accelerates flowering and increases elongation growth (Jung et al., 2016). In many cases, the process of growth can be differentiated by the response of plants to temperature (Went, 1953). Actually, only a few papers in which the temperature response is treated as a major problem can be mentioned (Yan and Hunt, 1999 and papers cited therein), but some recent publications have highlighted the role of temperature as a factor that regulates plant development, i.e. *via* phytohormones (Dockter et al., 2014; Jung et al., 2016).

Now, inspired by the Ansatz using Euler beta function *f*(*x*, α, β) for growth empirical data, we will show that the final effect of temperature and pH on plant growth is effectively similar, even though both of the triggers of these responses are apparently of a different nature.

Note that F is not free energy and G is not Gibbs energy throughout the article.

### Material and methods

#### Plant material

In order to check our equations, limited experiments were performed using three-day-old seedlings of *Zea mays* L cv. Cosmo. The maize seeds were soaked in tap water for 2h, and then sown in moist lignin. The seedlings were grown in an incubator in darkness at 27 ± 0.5°C for three days. Ten mm segments were cut from the plants 5 mm below the tip and first leaf was removed.

#### Simultaneous measurement of growth and acidification

The experiment was performed for artificial pond water (APW: 0.1 mM NaCl, 0.1 mM CaCl_2_, 1 mM KCl, pH 6.5 established by HCl or NaOH) conditions. Thirty ten-mm-long coleoptile segments were prepared and then placed in two identical sets (15) of the elongation-measuring apparatus in an aerated incubation medium. The volume of the incubation medium in the apparatus was 5 ml per glass tube (0.3 ml of the incubation medium per segment). Measurements of the growth and pH were performed for 12.5 hours and recorded every 15 minutes. The pH measurement was carried out using a CPI-501 pH-meter. Images of the segments were recorded using a CCD camera (Hama Webcam AC-150). Only one of the two tests in separate tubes was selected for analysis. The length of the coleoptile segments (initially 10-mm-long segments) was measured in the ImageJ program with the accuracy established at the ± 0.1 mm level. The relative elongation was calculated using the formula (*l*_t_ – *l*_0_)/*l*_0_ with *l* for “length” and *l*_0_ = *l*(*t* = *t*_0_) = 1 cm. Then, the relative volume was obtained by multiplying the length by the average cuticle segment cross-section area. The temperature during the experiment was 25 ± 0.5°C and was maintained using a water bath at a similar level. All manipulations were performed under dim green light.

#### Construction of a phenomenological model

Based on the “acid growth hypothesis” and relevant experimental data (Yan and Hunt, 1999 and Hager, 2003), we examined the pH/*T* (or μ/*T*) duality of acidic pH (or auxin-induced acidification) and temperature (*T*) for the growth of grass shoots in order to determine the model equation (EoS) for the extending primary wall of (living) plants.

In the literature, pH is defined as the decimal logarithm of the reciprocal of the hydrogen ion activity a_H+_ in a solution (pH = – log_10_ a_H+_ = log_10_ 1/a_H+_). The pH value is a logarithmic measure of H^+^-activity (the tendency of a solution to take H^+^) in aqueous solutions and defines their acidity or alkalinity; a_H+_ denotes the activity of hydronium ions in units of mol/l. The logarithmic pH scale ranges from 0 to 14 (neutral water has a pH equal to 7).

The temperatures at which most physiological processes normally occur in plants range from approximately 0°C to 45°C, which determines the physiological temperature scale (in Kelvin scale: [K] = [°C] + 273.15). The temperature responses of plants include all of the biological processes throughout the biochemical reactions (high Q_10_ factor). This is clearly visible in the relative rates of all of the development or growth processes of maize as a function of temperature, which is illustrated in Fig. 3 in Yan and Hunt (1999). For future applications, we propose that the ‘absolute’ temperature scale for plants [0–45°C], which corresponds to a [0, 1] interval after rescaling, be setup. Note, this is not an actual absolute temperature.

**Figure 3.**
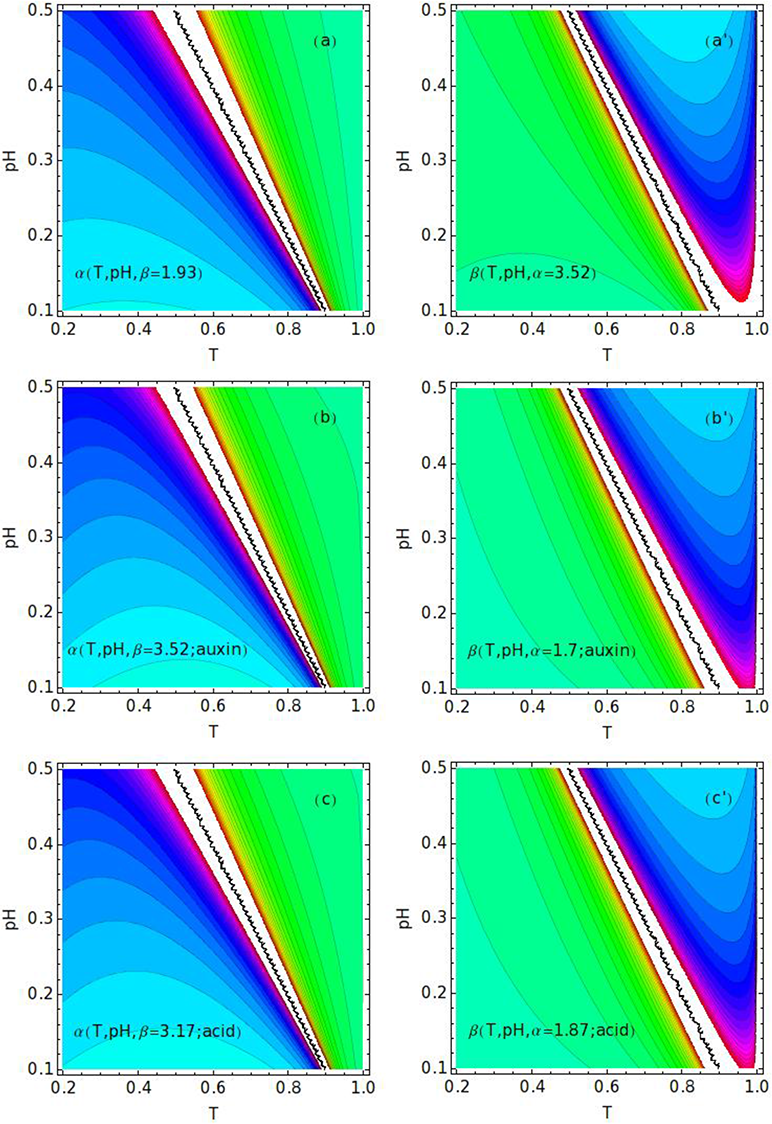
Solutions (state diagrams) for α and β exponents in the *T* and pH coordinates (rescaled). The lines of the (a, b, c) constant α- and the (a’, b’, c’) β-values in (*T*, pH) coordinates are indicated in the charts. Simulation data are presented in the figure legends. Green – negative values, blue – positive.

Which data have been adopted for the research and how those data have been handled is described below. For the analysis, we accepted the reliable published data, namely the pH plots that are presented in Fig. 2 in Hager (2003) and the temperature plots that are presented in Fig. 3 in Yan and Hunt (1999). These charts were first digitised by a GetData digitiser for reconstruction and then rescaled (normalised to [0, 1] interval) for both perpendicular axes (SI Figure 1), divided by a maximum value. The original temperature range [0, 45°C] was converted to a [0, 1] interval by dividing by 45. The same procedure was conducted on the pH scale [0, 14] in order to obtain the normalised [0, 1] interval (SI Figure 1), which was indispensable for the further calculations.

The pH plots that are presented in Fig. 2 in Hager (2003) as well as the temperature plots that are presented in Fig. 3 in Yan and Hunt (1999) look similar when mirror symmetry is applied. Providing that we perform a substitution *x* → (1 – *x*) and consider a function *f*(*x*) of variable *x* and of its reflection (1 − *x*), one might expect that they will look approximately the same after the correct scaling. These properties can be adequately described by the Euler beta density distribution (or beta Euler, for short). In this approach, we assume that *x* equals either pH or temperature (both rescaled to a [0, 1] interval), an assignment which makes sense since at a relatively low pH, which corresponds to a relatively high temperature in this unit-scaling, plant cells and organs grow (elongate) the fastest, which is clearly reproduced in SI Figure 1.

The data from SI Figure 1a were taken separately (“auxin” and “acid” growth; Hager, 2003) for the fitting procedure (SI Table 1 and SI Figure 2), while the temperature data (Yan and Hunt, 1999) from SI Figure 1b were first normalised and then fitted by beta Euler. Furthermore, when comparing the data from different experiments (Yan and Hunt, 1999; Hager, 2003), we observed that the *x* variable could, after rescaling, have the following meaning – either acidity (basicity) *x* ≡ pH or temperature *x* ≡ T (see SI Figure 1) with a characteristic beta function cut-off at *x* = 0 (SI Figure 1a and the inset) and at *x* = 1 (SI Figure 1b); the normalised scale we used is presented in SI Figure 1b (inset).

### Results

#### Derivation of the EoS for plants

The probability density function of the beta distribution (also called the Euler integral of the first kind (Polyanin and Chernoutsan, 2011) for 0 ≤ *x* ≤ 1 and shape parameters α and β > 0 is a power function of the variable x

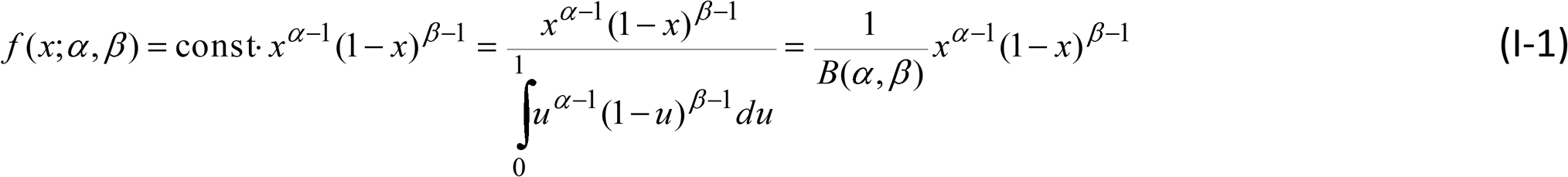

where *B* = *B*(α, β) is the normalisation constant. Based on the experimental observations as discussed in the previous sections and presented in SI Figure 1, the left side of equation (I-1), rewritten twice, first for *x* = pH, then *x* = *T*, can be merged into a single expression

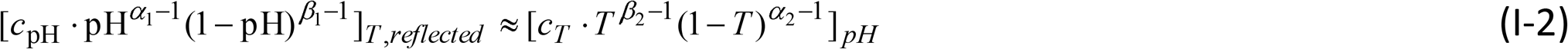

with a condition that *T* and pH are normalised and dimensionless; c_pH_ and c_T_ are constants normalising the maxima of the beta functions to 1, and “pH” is hereafter treated as a non-separable variable name. The lower indices “T” and “pH” in Eq. (I-2) should be read “at constant temperature” and “at constant pH”, respectively, which is valid for the preparation of strict experimental conditions (the use of a pH-stat apparatus as in Lüthen et al. (1990) to achieve a stable equilibrium pH seems to be indispensable; see also Hager, 2003). The appropriate (i.e. symmetric with respect to *x* = 0.5 line) values of T and pH should be taken into account in Eq. (I-2). The equality sign and the equalisation *α*_1_ = *α*_2_ =*α, β*_1_ = *β*_2_ =*β* as well as *c_T_ = c_pH_* in Eq. (I-2) seem to be justified for biological systems (see SI Figure 1.1) in the first order of approximation. Such a modified Eq. (I-2) will be considered further as a constraint on the two independent variables *T* and pH.

By defining the function *F* at constant turgor pressure *P* (isobaric conditions: δ*P* ≈ 0)

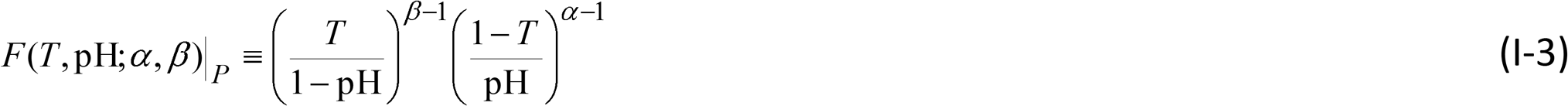

that is an analytic function of pH and *T* on the physically accessible interval (0, 1) for each variable (note that *F* is *not* the free energy of the system; here function *F* exclusively constructs a bridge between the physical and chemical features of the living system), and assuming adequate water uptake to fulfil this requirement, we determine from equation (I-2)

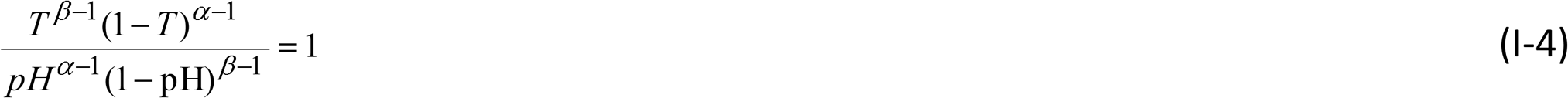

We note that Eq. (I-4) take, after transformation, the usual form of an equation of state. Hence, the constitutive EoS for living plant cells reads

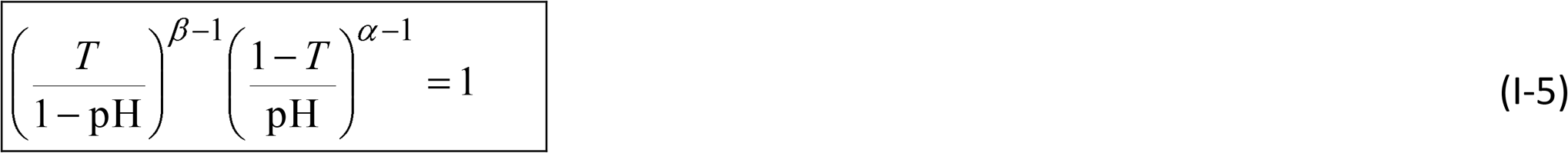

which has the elegant structure of a double power law. [Systems displaying power law behaviour in their various characteristics are abundant in Nature and the most impressive examples are found in biology (Racz, 2002)]. Note that *T* in equation I-5 is the temperature in Celsius rescaled to a [0, 1] interval as pH also is. Here, α and β are the shape exponents (SI Table 1 and SI Figure 2). The saddle-point type of the solution that is presented in Fig. 1 represents a concave surface in the form of a contour plot, which is roughly accurate at low pH and moderate temperatures. Equation (I-5) becomes increasingly inaccurate (divergent) at very low or very high pH values (close to zero or one in the normalised scale, which we use throughout the article), even though such excessive conditions, which constitute unacceptable extremes for life to come to existence, are excluded by Nature. The isotherms of Eq. (I-3) are presented in Fig. 2a, while the lines of constant pH are presented in Fig. 2b. Note that the use of the term “equation of state” can be questionable, since it does not represent the relation between pH and *T* in the system from a thermodynamic point of view. Therefore, we use EoS instead.

Even though we (apparently) launched our derivation from the “acid growth theory”, a closer look at the ‘initial conditions’ revealed that, in fact, it was established on the raw experimental data of pH and *T* dependent growth (SI Figure 1). For that reason, Eq. (I-5) is independent of whether the acid growth theory applies or not, and therefore, seems universal, at least for tip-growing grass shoots. Note, Eq. (I-5) is not an evolution (growth) equation, but a system (cell wall) property equation. The solutions of Eq. (I-5) for α and β in the form of contour plots are presented in Figure 3.

The essential relations are property relations and for that reason they are independent of the type of process. In other words, they are valid for any substance (here: the primary wall of a given plant species) that goes through any process (mode of extension). Yet, in our case, we should be aware of the empirical origin of Eq. (I-5) and remember the fact that a fundamental variable of the system is the chemical potential μ at a constant pressure and temperature (here: μ = μ_H+_(*T*) for H^+^ ions). By taking the total derivative, where the implicit μ_H+_(*T*) dependence is taken into account, we get

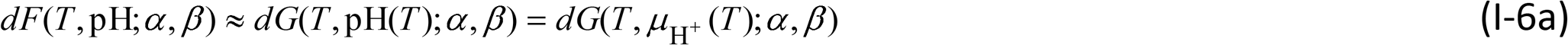

where

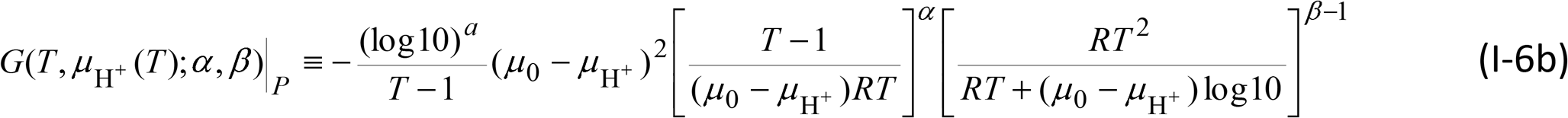

G_P_(*T*, μ) is another function (which should *not* be confused with the Gibbs energy) although expressed by intensive, microscopic variables. In approximation, since pH is usually measured in isothermal conditions (SI Figure 1a), we can temporarily suspend the pH(T) dependence in equation (I-6a) and consider pH and *T* as intensive (non-additive) variables. Note, that Eqs. (I-3) and (I-6) do not conclude about the free energy of the cell wall – they only describe the mathematical relations between pH, *T* and μ_H+_ variables.

Equation (I-5) is the EoS that describes the plant cell (wall) properties, where the SFs are the scaled temperature and pH of the expanding cell volume. Here, pH, which is intimately connected with the acid growth hypothesis even though it is not an intensive variable in the usual sense, introduces a direct link between the physical variables that represent the state of the system (growing cell or tissue) and the biological response (*via* pH altering – H^+^-ATPase), which results in growth. Clearly, equation (I-5) can be further verified by the diverse experiments that have been conducted for many species in order to determine the characteristic triads (α, β, Ω_S_) that belong to different species or families (classes) taxonomically, or that are simply controlled by growth factors (growth stimulators such as auxin, fusicoccin or inhibitors such as CdCl_2_). However, the real test of the utility of Eq. (I-5) would include the predictive outputs for any perturbations that could then be subject to experimental validation. From the dyads (α, β), Eq. (I-5) should also be able to connect with either the underlying physiological processes or with their molecular drivers. In particular, whether the underlying molecular mechanism is identical or different should be reflected in the critical exponents (α and β pair should be identical for the same molecular mechanism and dissimilar for diverse mechanism). Despite the fact that F is not a generalised homogeneous function, it can also be used to check for a kind of “universality hypothesis” (Stanley, 1971) and to validate the evolutionary paradigm with genuine numbers.

#### Calculating critical exponents

Analytical analysis must lead to predictions that can be used as a Litmus test in order to cut through the myriad of conceptual models that have been published in the biophysical community. In plant science, given that the calculation of critical exponents can be connected with the optimum growth, it can be helpful to identify the molecular mechanism that is responsible for growth (in principle, similar exponents should be connected with the same mechanism). In physics, phase transitions occur in the thermodynamic limit at a certain temperature, which is called the critical temperature, *T*_c_, where the entire system is correlated (the radius of coherence ξ becomes infinite at *T* = *T*_c_). We want to describe the behaviour of the function *F*(pH, *T*), which is expressed in terms of a double power law by equation (I-3) that is close to the critical (optimum) temperature and, specifically in our case for practical reasons, also about the critical (optimum) pH = pH_c_. Critical exponents describe the behaviour of physical quantities near continuous phase transitions. It is believed, though not proven, that they are universal, i.e. they do not depend on the details of the system. *Per analogia*, since growth (such as cell wall extension in pollens or the elongation growth of the shoots of grasses, coleoptiles or hypocotyls) can be imagined as advancing over the course of time, a series of non-equilibrium STs (see Discussion), the critical exponents can be calculated.

Let us introduce the dimensionless control parameter (τ) for a reduced temperature

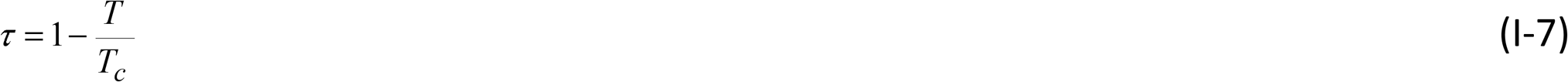

and similarly (π), for a reduced pH

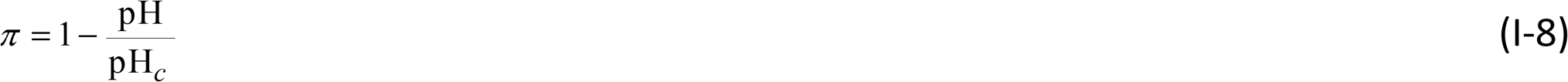

which are both zero at the transition, and calculate the critical exponents. It is important to keep in mind that (in physics) critical exponents represent the asymptotic behaviour at a phase transition, thus offering unique information about how the system (here: the polymer built cell wall) approaches a critical point. A critical point is defined as a point at which ξ = ∞ so in this sense *T* = *T*_c_ and pH = pH_c_ are critical points of Eq. (I-5). We believe that this property can help to determine the peculiar microscopic mechanism(s) that allows for wall extension (mode of extension) and growth in future research. The information that is gained from this asymptotic behaviour may advance our present knowledge of these processes and their mechanisms and help to establish the prevailing one.

By substituting equations (I-7) and (I-8) into equation (I-3), we can calculate the critical exponents (when *T → T_c_* and pH → pH*_c_*) from the definition (Stanley, 1971), which will give us

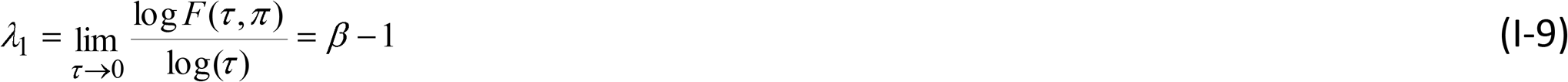

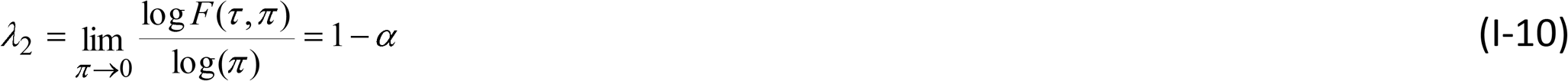

and

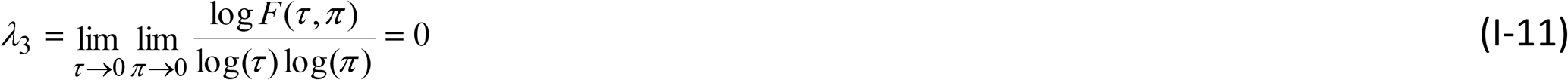

Here λ1 = β – 1 and λ_2_ = 1 – α are the critical exponents (λ_3_ = 0). The above equations result in two power relations that are valid in the immediate vicinity of the critical points

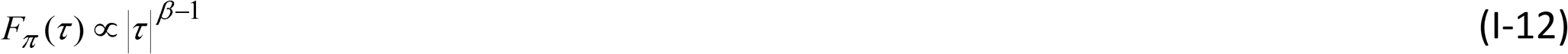

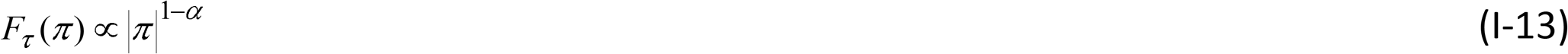

where lower indices denote constant magnitudes. Equations (I-12) and (I-13) represent the asymptotic behaviour of the function F(τ) as τ → 0 or F(π) as π → 0. In fact, we can observe a singular behaviour at *T* = *T*_c_ (Fig. 4a) and pH = pH_c_ (Fig. 4b). At this bi-critical point, the following relation holds: α + β = 2.

**Figure 4.**
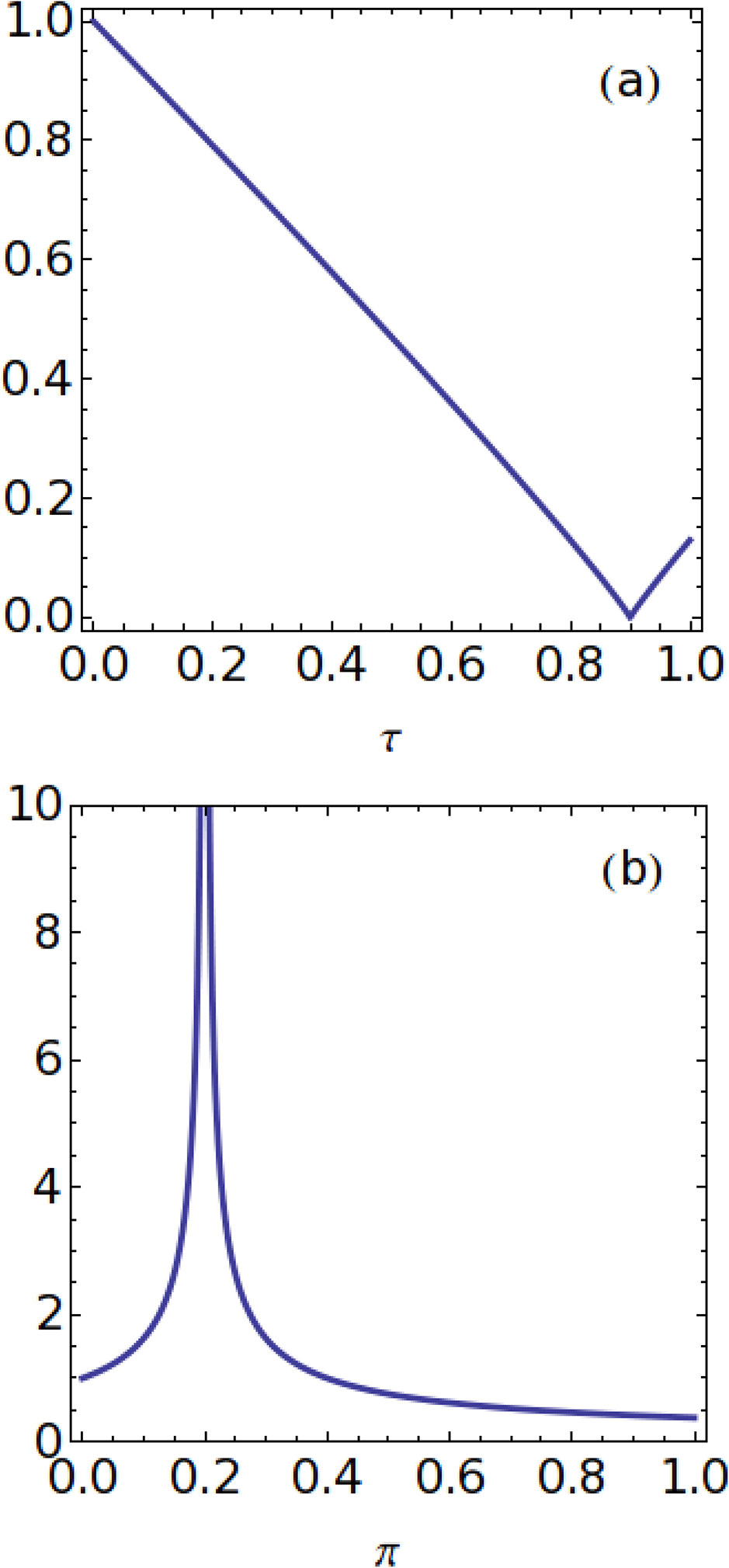
Critical behaviour at the STs that take place in the cell wall (pH, *T* – variables). (a) A “kink” (discontinuity in the first derivative) at the critical temperature *T* = *T*_c_. (b) a “log-divergent” (“λ” – type) solution at the critical pH = pH_c_. Control parameters: reduced temperature (τ) and reduced pH (π). Simulation parameters: (a) β = 1.93 and (b) α = 1.7 (SI Table 1).

Retaining τ for a reduced temperature, let us introduce another – microscopic – control parameter, μ, for the chemical potential

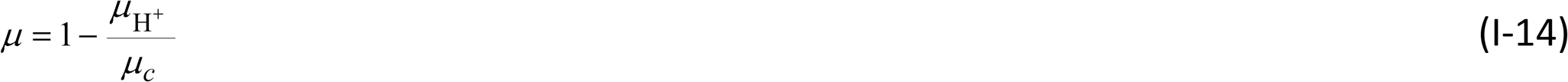

Substituting τ and equation (I-14) into equation (I-6b) for G(T, μ) and assuming the reference potential μ_0_ = 0, we can calculate the critical exponents (when *T → T_c_* and 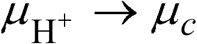) to get

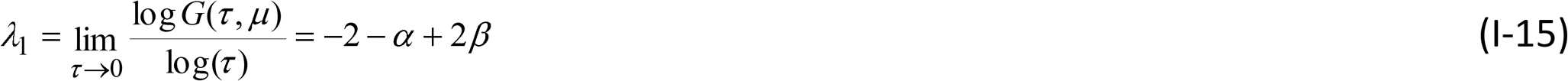

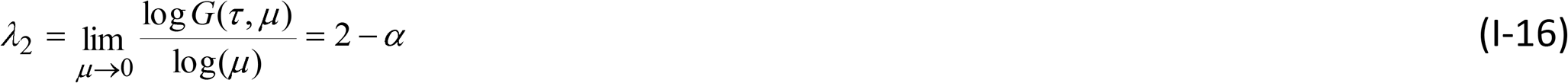

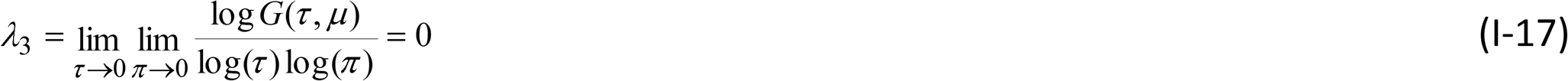

The above limits result in two more power relations, which are valid in the immediate vicinity of the critical points

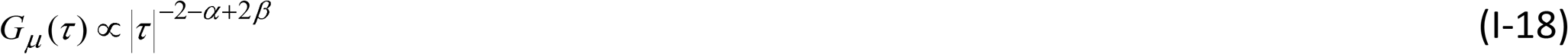

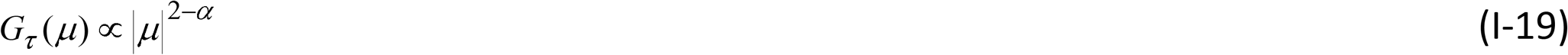

Equations (I-18) and (I-19) represent the asymptotic behaviour of the function G(τ, μ) as τ → 0 or G(τ, μ) as *μ* → 0. At the microscopic level, we observe a singular behaviour at *T* = *T*_c_ (Fig. 5a) and μ = μ_c_ (Fig. 5b-c).

**Figure 5.**
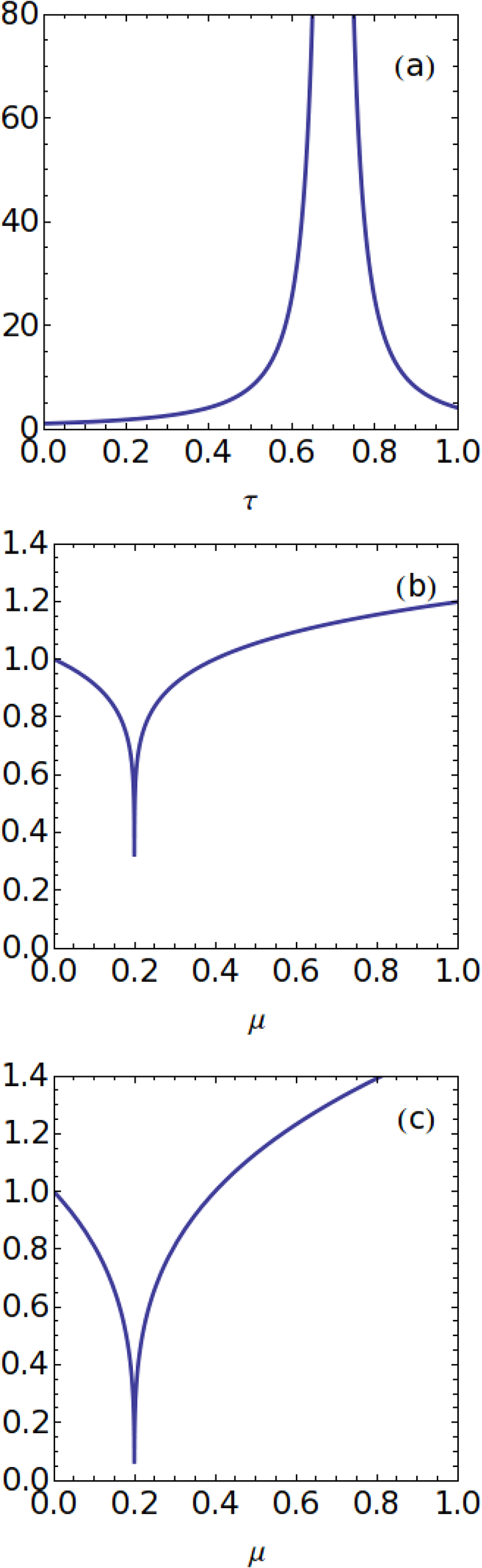
Critical behaviour at the STs that take place in the cell wall (for μ, *T* – variables). (a) A “log-divergent” (“λ” – type) solution at *T* = *T*_c_. (b – c) Solution at the critical value of the chemical potential μ = μ_c_; (b) “acid growth”, (c) “auxin growth” (SI Table 1). Control parameters: reduced temperature (τ) and chemical potential (μ). Simulation parameters: (a) α = 3.52, β = 1.93; (b) α = 1.87, β = 3.17; (c) α = 1.7 and β = 3.52 (SI Table 1).

For the bi-critical point, the exact relation β = 2 holds (compare with SI Table 1 and SI Figure 2), thus leaving us with the only free parameter (α) of the theory. Note the broad peak at the critical temperature (indicating that optimal growth occurs within a certain temperature range) and a sharp, more pointed peak for the critical chemical potential (a ‘single value’ that corresponds to the optimum growth). In this representation, growth can be treated as a series of subsequent STs that take place in the cell wall (see Part II). It looks as though a cardinal quantity that is decisive for growth in this particular approach is the chemical potential. During the period of the evolution of the extending cell wall, the chemical potential could have formed asymmetrical ratchet mechanism. Oscillating biochemical reactions, which are common in cell dynamics, may be closely related to the emergence of the phenomenon of life itself (Martin et al., 2009). In this context, the relations for the chemical potential may also belong to the fundamental dynamic constraints for the origin of life.

Complex systems such as the cell wall of a growing plant are solid and liquid-like at the same time. At the molecular level, they are both ordered and disordered (allowing for the transition between these states). There are obviously complex interactions between the model parameters and inputs from many other kinds of inputs. However, the EoS describes the properties of the wall and unknown complex interactions in a general way. Whether the wall structure is ordered or disordered, universal features can be identified using simple scaling laws.

### Discussion

Erwin Schrödinger (1944) may have been the first to consider the thermodynamic constraints within which life evolves, thereby raising fundamental questions about organism evolution and development. Here, we attempted to present a more modest description in terms of the EoS and methods such as the calculations of critical exponents. More precisely, this work seeks to model the relationship between plant growth and temperature (or pH) in terms of the EoS, thereby describing growth optima as functions of the ‘state’ variables and critical exponents. Apparently, there is already a simple and compelling explanation for the growth optima of plant growth from the biological point of view. Growth is mediated both directly and indirectly by enzymes and increases broadly in line with the frequently observed effect of temperature on enzyme activities (Berg et al., 2002). However, once temperatures are sufficient to denature proteins, enzyme activities rapidly decrease and plasma membranes and some other biological structures also experience damage (compare SI Figure 1b at a high temperature).

#### Sensing structural transitions

A magnitude that infinitesimal changes result in symmetry change always exists in physics in continuous phase transitions (Landau and Lifshitz, 1980). However, structural changes such as the ones considered here usually take place in non-equilibrium conditions (Racz, 2002). Seemingly, such (order/disorder) symmetry changes (De Gennes, 1991) in biological systems may be connected with a mechanism (Rojas et al., 2011) by which chemically mediated deposition causes the turnover of cell wall cross-links, thereby facilitating mechanical deformation. It may also reflect the pectate structure and distortion that was suggested by Boyer (2009) (Fig. 5c, d) in *Chara* cell walls by placing wall polymers in tension and making the load-bearing bonds susceptible to calcium loss and allowing polymer slippage that irreversibly deforms the wall. Proseus and Boyer (2007) suggested that a ladder-like structure would be susceptible to distortion and that the distortion would increase the distance between adjacent galacturonic residues (Fig. 5d in Boyer, 2009), and therefore, the bonds may lengthen and thus weaken and decrease their affinity for Ca^2+^. Dissociation may then occur, thus allowing an irreversible turgor-dependent expansion (see also Fig. 4 in Pietruszka, 2013; this is also applicable for non-isochronous growth in pollen tubes). The direction of the maximal expansion rate is usually regulated by the direction of the net alignment among cellulose microfibrils, which overcomes the prevailing stress anisotropy (Baskin, 2005). However, the organisation of cellulose microfibrils depends on the organisation of the cortical microtubules (Paredez et al., 2006), which is modified by many different factors such as mechanical stimuli, light or hormones (Fishel and Dixit, 2013). These modifications are regulated *via* DELLA proteins (Locascio et al., 2013) and/or arabinogalactan proteins (Willats and Knox, 1996; Marzec et al., 2015b). The transient changes in the wall composition and the deformation of bonds may be a hallmark of symmetry change and a kind of ST (orchestrated instability) that is taking place in the cell wall.

The cell wall is supplied with the precursors of polysaccharides and also some proteins through exocytosis. This process is a form of active transport in which a cell transports molecules out of the cytoplasm into the extracellular matrix or intracellular space by expelling them in an energy-using process. Endocytosis is its counterpart. The loaded vesicles are transported by kinesin through the microtubules into the plus end while x toward the minus ends the, and ATP is utilised in both the binding and unbinding processes of the motor heads. Besides microtubules, the actin cytoskeleton represents the most demanding energy-using process. In fact, in plant cells, both exocytosis and endocytosis are F-actin dependent processes that are related to cell wall assembly and maintenance; moreover, there is strong evidence that plant cell elongation itself also depends on motors actin cytoskeleton rearrangement, see a series of studies by Baluška et al. (1997, 2001 and 2002) and Wojtaszek et al. (2007). Indeed, these energy-consuming processes are apparently not included in the EoS.

A situation that was already pointed out in the Introduction in which the protons that are excreted into the wall compartment are directly responsible for the wall-loosening processes through the hydrolysis of covalent bonds or the disruption of non-covalent bonds may also be a signature for the symmetry change and STs that occur in the wall compartment of the growing cell. Though the underlying microscopic mechanisms are not well recognised as yet (and as a result the ‘order parameters’ are difficult to identify), a pH-driven “lambda” ST may be attributed to the maximum activity of PM H^+^-ATPase, while temperature-driven STs in the cell wall can be directly related to the maximum elevation of the effective diffusion rate (*k*_2_ coefficient in Pietruszka, 2012) – we considered primary (diffusive) growth throughout this article. As an aside, from the calculation of cross-correlations, we may draw further conclusions that are connected with the biochemical picture of the acid growth theory. The results presented in SI Figure 3 is a clear characteristic that temperature-induced growth and auxin-induced acidification (PM H^+^-ATPase) growth correlate strikingly well (there is good experimental evidence that a higher temperature induces auxin biosynthesis in some plants). However, the convolution of acidic pH growth and temperature growth is less shifted away from zero (lag) and is even more pronounced (see also recent outcomes in Pietruszka and Haduch-Sendecka (2016) as well as Olszewska et al. (2017) for cell wall pH (proton efflux rate) and growth rate, which are directly co-regulated in growing shoot tissue, thus strongly supporting the empirical foundations of EoS). This strict quantitative result may contribute to the acid growth hypothesis that was developed by plant physiologists and make a contribution to resolving the problems that occur in the long-lasting discussion. However, a problem still arises. It is unclear to what degree Eq. (I-5) applies universally as the published raw data that constitute the experimental basis are only for grass shoot elongation growth. The long-standing acid growth theory postulates that auxin triggers apoplast acidification, thereby activating the cell wall-loosening enzymes that enable cell expansion in shoots. This model remains greatly debated in roots. However, recent results (Barbez et al., 2017) in investigating apoplastic pH at a cellular resolution show the potential applicability of Eq. (I-5) for roots.

We now show that our findings may be embedded in the evolutionary context that is connected with the migration of plants away from the Equator (changes in latitudes) as the climate changed as well as their adaptation to the spatial distribution of pH in the soil as a substitute for a high temperature. As an example that exhibits the capabilities of equation (I-5), let us consider the following. The dominant factors that control pH on the European scale are geology (crystalline bedrock) in combination with climate (temperature and precipitation) as was summarised in the GEMAS project account (Fabian et al., 2014). The GEMAS pH maps primarily reflect the natural site conditions on the European scale, whilst any anthropogenic impact is hardly detectable. The authors state that the results provide a unique set of homogeneous and spatially representative soil pH data for the European continent (note Fig. 5, ibid.). In this context, the EoS that is expressed by equation (I-5) may also have an evolutionary dimension when considering the spatial distribution of pH data as affected by latitude (angular distance from the Equator) elevation on the globe, which is associated with climate reflection (ibid.). pH is strongly influenced by climate and substrate – the pH of the agricultural soils in southern Europe is one pH unit higher than that in northern Europe. See also Fig. 4 in Fabian et al. (2013) in which the pH (CaCl2) of soil samples that are grouped by European climate zones are measured (descending from pH ≈ 7 (Mediterranean), pH ≈ 6 (temperate) through pH ≈ 5.25 (boreal) to pH ≈ 4.80 (sub-polar)). This fact may be reinterpreted in terms of the pH-temperature duality that is considered in this work. When the migration of plants away from the Equator occurred, lower pH values further to the North might have acted in a similar way as high temperatures on the growth processes in the tropic zones. This situation could have led to energetically favourable mechanisms (adaptation) such as the amplified ATP-powered H^+^ extrusion into the extending wall of plant cells and the secondary acidification of the environment (released protons decrease pH in the incubation medium), irrespective of the initial pH level, which is the main foundation for the acid growth hypothesis. Apparently, the evolutionary aspect of equation (I-5), which suggests a possible link with self-consistent adaptation processes (“The individual organism is not computed, or decoded; it is negotiated” (Walsh, 2010), as one of its first applications, seems difficult to underestimate, and the potentially predicative power of EoS looks remarkable considering the modest input we made by deriving equation (I-5). In this perspective, the pH-response could potentially provide a plant with an adaptive advantage under unfavourable climate conditions. Together with the quite recent discovery of the function of phytochromes as thermosensors (Jung et al., 2016), the EoS could be useful in breeding crops that are resilient to thermal stress and climate change.

#### Perspectives

The EoS for a biological system has not been reported until this study. To further verify the EoS and its implications, we note that based on the results that were obtained (calculated ‘critical exponents’), a kind of “universality hypothesis” (Stanley, 1971) can be tested experimentally in order to extract “classes of universality” for plant species (Ω_S_), thereby substantiating the taxonomic divisions in the plant kingdom *via* quantitative measures (Table 1). This issue, which we may call “extended plant taxonomy”, demands further experimentation and is beyond the scope of this theoretical work.

**Table 1.**
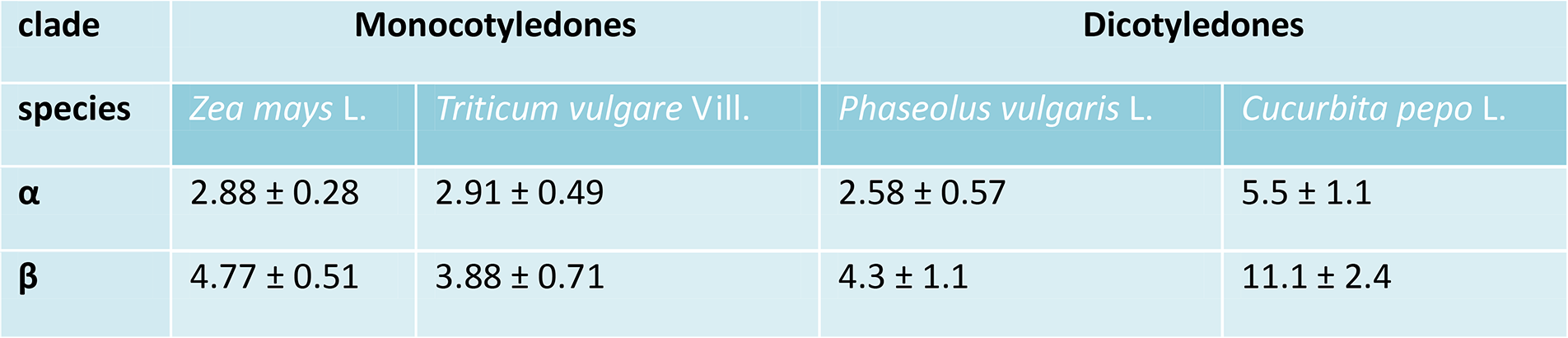
Preliminary data for α and β for maize (*Zea mays* L.) and wheat (*Triticum vulgare* Vill.), which belong to the same family *Poaceae* but to different subfamilies: *Panicoideae* and *Pooideae*, respectively; the calculated values for α and β are similar. The common bean (*Phaseolus vulgaris* L.) and cucurbita (*Cucurbita pepo* L.) belong to the same clade *Eudicots* but to different orders: *Faboles* and *Cucurbitales*, respectively; the calculated values for α and β differ significantly. The calculation that is based on Fig. 1 in Lewicka and Pietruszka (2008) and Eq. (I-5). Table courtesy of my PhD student A. Haduch-Sendecka.

| clade | Monocotyledones |  | Dicotyledones |  |
| --- | --- | --- | --- | --- |
| species | *Zea mays* L. | *Triticum vulgare* Vill. | *Phaseolus vulgaris* L. | *Cucurbita pepo* L. |
| $\alpha$ | $2.88 \pm 0.28$ | $2.91 \pm 0.49$ | $2.58 \pm 0.57$ | $5.5 \pm 1.1$ |
| $\beta$ | $4.77 \pm 0.51$ | $3.88 \pm 0.71$ | $4.3 \pm 1.1$ | $11.1 \pm 2.4$ |

It seems the EoS may explain a number of phenomena and is powerful in its simplicity. It positions itself at the brink of biology and nano-materials. We believe it may be helpful in identifying the processes of the self-assembly of wall polymers. In agricultural implementations, any departures from the optimum growth may simply be corrected by appropriately adjusting pH or T.

The finding of the EoS for plants adds an important dimension in the biophysical search for a better explanation of growth-related phenomena in a coherent way through its applicability to the visco-elastic (or visco-plastic) monotonically ascending and asymptotically saturated (Pietruszka, 2012) cell wall extension that is usually observed, as well as to pollen tube oscillatory growth (Part II). The underlying biochemical foundation, which is expressed in the form of the acid growth hypothesis, could also help to understand the growth conditions that are ubiquitous in biological systems through coupling to EoS. In this respect, our method is accompanied by its partial validation – its application to the important biological question concerning the acid growth hypothesis. In the above context, it is not surprising that the growth of the cell wall in plants can be thought of as a time-driven series of STs (*via* the breaking of polymer bonds or other mechanisms that were mentioned earlier), although they are ultimately connected with the physical conditions and chemistry by the EoS (equations (I-5) or (I-6b)), which are solved at subsequent time instants. Since non-equilibrium stationary states, which always carry some flux (of energy, particle, momentum, etc.), are achieved after discrete intervals of time for single cells such as pollen tubes, such a (chemical potential ratchet) mechanism can lead to a kind of ‘leap’ in the macroscopically (of μm scale) observed growth oscillations (Geitmann and Cresti, 1998; Hepler et al., 2001; Zonia and Munnik, 2007). Similar mechanisms were observed in multicellular models for plant cell differentiation such as rhizodermis (Marzec et al., 2013a) where the growth of root hair tubes occurs by leaps and bounds (Vazquez et al., 2014). This phenomenon may be treated as the final result of the subsequent STs that take place in the cell wall.

On the other hand, spontaneous bond polarisation/breaking may be synchronised through Pascal’s principle by pressure P or pressure fluctuations δP (Pietruszka and Haduch-Sendecka, 2015a), thereby acting as a long-range (of the order of ξ) messenger at the state change (the radius of coherence *ξ* → ∞ at a Γ-interface (Pietruszka, 2013)). When the entire system is correlated and the stress-strain relations are fulfilled (ibid.), the simultaneous extension of the cell wall at the sub-apical region may correspond to pollen tube oscillation(s). Hence, an uncorrelated or weakly correlated extension would result in the bending that is observed in the directional extension of the growing tube (ibid.). In multi-cell systems (tissues) that have a higher organisation (coleoptiles or hypocotyls in non-meristematic zones), the EoS for plants will manifest itself by the emergent action of an acidic incubation medium or will be induced by endogenous auxin acidification as is presumed in the acid growth theory. In this aspect, the EoS may serve as a new tool for the further investigation and verification of claims about the acid growth hypothesis (see Barbez et al., 2017 for recent results). This should greatly facilitate the analysis of auxin-mediated cell elongation as well as provide insight into the environmental regulation of the auxin metabolism.

It is important to note that “Cellular growth in plants is constrained by cell walls; hence, loosening these structures is required for growth. The longstanding acid growth theory links auxin signalling, apoplastic pH homeostasis, and cellular expansion, providing a conceptual framework for cell expansion in plant shoots. Intriguingly, this model remains heavily debated for roots” (Barbez et al., 2017). The authors’ findings permitted the complex involvement of auxin in root apoplastic pH homeostasis to be elucidated, which is important for root cell expansion, and shed light on the acid growth mechanism in roots. This is, in turn, the outcome of our model equations, which are apparently applicable for elongating roots (root hair), shoots or even a single plant cell (pollen tube).

Moreover, by using the EoS, it will be possible to answer the question about role of the crosstalk between auxin and other hormones such as strigolactones (Marzec and Muszynska, 2015), in growth of plant cells. It also seemingly delivers a narrative that provides a biophysical context for understanding the evolution of the apoplast, thereby “uncovering hidden treasures” in the as yet unscripted biophysical control systems in plants.

### Summary

This section takes a physical approach to plant growth and its aim is to simplify the universal features that may explain the observations. We have fitted a simple algebraic function (a beta Euler function) to some growth rate data, thus capturing the variations with respect to temperature and pH, which show a surprisingly conjugate nature.

Cross-disciplinary research at the interface between the physical and life sciences was also accomplished in this part. We first considered the Euler beta function enigma, which is the duality of acidic pH (or auxin-induced acidification) and temperature, in order to obtain the EoS for a living system. We started from the striking similarity (mirror symmetry) between the elongation growth of the grass shoots that were incubated in different pH environments or auxin-induced elongation in water (or a neutral buffer) with the respective growth at different temperatures. We based these on the hypothesis that the effect of temperature on elongation growth is effectively equivalent for the relative growth increments such as those that are caused by a change of pH or endogenous auxin-induced acidification in the incubation medium. In order to resolve this ambiguity, we first used the beta function for both dependences, which were normalised prior to the comparison, in order to obtain the beta function shape parameters α and β. It turned out that even without referring to the biochemical underpinning, we might have concluded that the acidic conditions of the incubation medium or auxin-induced acidification and environmental temperature can act interchangeably, at least when they are considered at a phenomenological (mechanistic) level, i.e. from the effectiveness of the point of view of plant growth. The numerically verified high accuracy of these complementary representations allowed us, among others, to treat this dual approach as a new apparatus for predicting the outcomes in different growth conditions, which is especially useful in changing climate environments. We presumed that by applying a beta distribution and continuously changing its character with the values of the shape parameters, our findings might be related to the evolutionary context that is connected with the migration of plants away from the Equator as the climate changed and their adaptation to the spatial distribution of pH in the soil as a substitute for high temperatures. The EoS may be also helpful in delivering analytical solutions (the respective (α, β, Ω_S_) – triads) for the assisted migration of plant species (Vitt et al., 2010) that are at the risk of extinction in the face of rapid climate change. In this application, the EoS tool can constitute a complementary method for genetic modifications in the assisted migration processes by permitting the best species to be selected for such an introduction from those having a similar (to within experimental error) triad.

In physics, it is believed (Fisher, 1966; Griffiths, 1970) that the critical exponents are “universal”, i.e. independent of the details of the Hamiltonian (energy function) that describes a system. The implications of this, however, are far reaching. One could take a realistic and complicated Hamiltonian, ‘strip’ it to a highly idealised Hamiltonian and still obtain precisely the same critical exponents. For instance, on these grounds, it is believed that carbon dioxide, xenon and the three-dimensional Ising model should all have the same critical exponents (Baxter, 1989) and this appears to be the case (Hocken and Moldover, 1976). In this context, since growth phenomenology (non-equilibrium orderings) and the STs approach have converged to a form an EoS for plants, the potential role of the critical exponents in discriminating the different modes of the wall extension (Breidwood et al., 2014 for review) of growing cells or tissues seems promising in opening new avenues of research.

## PART II Equation of Evolution for Plants

### Introduction

In this section, we pose the question of whether the cell volume extension or elongation growth of plants (represented by expanding volume V) can be predicted by pH, T and P.

Modulation of the mechanical properties during expansive growth is a ‘hot topic’ for the plant cell growth community, e.g. recently in Bidhendi and Geitmann (2015), Boudon et al. (2015), Rojas et al. (2011) and Barbacci et al. (2013). Plant cell expansion, in turn, is controlled by the balance between the intracellular turgor pressure, cell wall synthesis and stress relaxation. Several mainstream explanations of the mechanical aspects of cell growth have been considered in publications, beginning with the cell wall model (Holdaway-Clarke and Hepler, 2003) through the instability model (Wei and Lintilhac, 2007) to the hydrodynamic model (Zonia and Munnik, 2007). In this section, we propose that these apparently conflicting visions of cell wall extension and growth may be consolidated in a unifying chemical potential based explanation, which hopefully converges in a coherent picture of this somewhat elusive topic.

The last few years have witnessed significant progress in uncovering the molecular basis that underlies plant cell elongation and expansion. In many cases, the first step in cell growth is related to a restriction of the symplasmic communication between neighbouring cells, which permits a specialisation programme that is specific for the individual cell to begin (Marzec and Kurczynska, 2014). This isolation enables cells from multicellular organisms to be considered to be individual cells. Afterwards, different signalling and developmental mechanisms are run in order to induce either the expansion of the entire cell or localised elongation. The roles of different protein classes have been described based on the analysis of the growth of pollen or root hair tubes, which are the models that are used in investigations of tip-growth (Higashiyama et al., 2015; Marzec et al. 2015a). Among them, the function of expansins, the proteins that are involved in plant cell wall loosening, which is related to cell wall acidification, has been confirmed in both monocot and dicot species. Analysis of different mutants in the genes encoding expansins confirmed their role based on the analysis of plant phenotypes (Cosgrove, 2015). Nevertheless, a model that permits the cell status of mutant and wild-type plants to be compared might prove useful for further investigations.

According to the acid growth hypothesis, auxin plays a pivotal role in the growth of plant cells (Hager, 2003; Lüthen, 2015 for review). However, recent reports have indicated that a new class of plant hormones, strigolactones (SLs), cooperate with auxin in many developmental processes such as shoot and root branching under normal and stress conditions (Marzec et al., 2013b). Additionally, new functions of SLs, which indicate that those hormones may play a key role in the regulation of plant growth and development, have recently been proposed (Marzec and Muszynska, 2015). Until now, most data has indicated that SLs influence auxin transport *via* the distribution of PIN proteins (Shinohara et al., 2013), but it has also been proposed that SLs influence secondary growth in plants (Agusti et al., 2011) and cell elongation (Hu et al., 2010; Koltai et al., 2010). Mathematical formulas that permit cell growth under different treatments to be measured and described may shed new light on the hormonal regulation of plant cell elongation.

From the physical point of view, the basic ingredients of an expanding plant cell are the cell wall, plasma membrane and cytoplasm. Together, they form a system that can be described as a two-compartment structure with the plasma membrane being treated as the interface between the cytoplasm and the wall. On the other hand, the intensive quantities can be attributed to the system of volume V, namely pressure (P), and the chemical potential (μ), the latter of which is directly related to a measurable quantity – pH. The problem of the volumetric growth of plant cells, which lies at the intersection of physics, biochemistry and composite materials of a nanometric scale, is inherently difficult to capture in a single theoretical frame (Braidwood et al., 2014). Therefore, the goal was to deconstruct all of the above-mentioned approaches in order to assemble a model that is based on physical principles that are also able to admit other perspectives. We believe that the EoS that was proposed above has the ability to do this, although its relevance to many of the growth details may be beyond the scope of this conjecture.

#### Chemical wall loosening approach

It is believed that the cell wall has active mechanisms for self regulation (Hepler and Winship, 2010). The cell wall is formed through the activity of the cytoplasm and maintains a close relationship with it. Communication occurs between the cell wall and the cytoplasm and the plasma membrane occupies a pivotal position in transmitting particles and information. In this model, when applied to pollen tube growth, cell growth is ultimately dependent on the biochemical modification of the wall at the apical part where exocytosis is believed to occur (Holdaway-Clarke and Hepler, 2003). Cell wall loosening then allows turgor-induced stretching, thereby pulling the flow of water into the cell (down its potential gradient). During oscillatory growth, enzymes that interfere with wall loosening are believed to be periodically activated or inhibited, which is supposed to drive oscillations in growth and the growth rate (as commented by Zonia, 2010). In this approach, the passive role of turgor pressure is stressed.

#### Loss of stability model

A new look at the physics of cell wall behaviour during plant cell growth, which focuses on the purely mechanical aspects of extension growth such as wall stresses, strains and cell geometry, was investigated within the framework of a model that was based on the Eulerian concept of instability (Wei and Lintilhac, 2003; 2007). It was demonstrated that loss of stability (LOS) and consequent growth are the inevitable result of the increasing internal pressure in a cylindrical cell (such as the internodal cylindrical cells of intact *Chara corralina* plants) once a critical level of pressure-induced stress (P_c_) is reached. The predictions of this approach were confirmed by direct measurements (ibid.). This (LOS) model apparently challenged the visco-elastic or creep-based models (Ray et al., 1972; Cosgrove, 1993; Schopfer, 2006), which were designated as chemical wall loosening and was questioned by Schopfer (2008), who asserted that stress relaxation can be attributed to chemorheological changes in the load-bearing polymer network that permit the plastic deformation of wall dimensions. In his words, the “growth of turgid cells is initiated and maintained by chemical modifications of the cell wall (wall loosening) followed by mechanical stress relaxation generating a driving force for osmotic water uptake”. In response to Schopfer’s letter, Wei and Lintilhac (2008) claimed that LOS did not challenge the principles of osmotic water relations or biochemically mediated wall loosening and emphasised that plant cells are both osmometers and pressure vessels, and therefore, both facts should be accommodated in any meaningful model. However, the problem of the water potential difference ΔΨ = Ψ_o_ – Ψ_i_ of a growing cell and a “mysterious pump” if pressures must rise to drive LOS behaviour still remained. Regardless of whether the new wall synthesis is homogeneous or patchy (the cell wall can always be conceptually shrunk to a patch), it will affect cell behaviour by modifying the turgor pressure P in short time scale oscillations or pseudo-chaotic fluctuations δP (Pietruszka and Haduch-Sendecka, 2015a), which are specifically enhanced by the flow of the osmotic pressure through the entire cell volume (beyond the apex as well).

#### Hydrodynamic model

Recent works have revealed an important role for osmotic pressure and the hydraulic features of the system in driving cell shape restructuring and growth (Proseus et al., 2000; Proseus and Boyer, 2006; Zonia et al., 2006, 2010; Haduch-Sendecka et al., 2014). This concept was established by works that revealed an important role for hydrodynamics in pollen tube growth. The main message that hydrodynamics have the potential to integrate and synchronise the function of the broader signalling network (in pollen tubes) was expressed by Zonia (2010). To be accurate, Pascal’s principle states that pressure exerted anywhere in a confined incompressible fluid (cell sap) is transmitted in all directions equally throughout the fluid in such a way that the pressure variations (initial differences) remain the same. Indeed, according to Pascal’s principle, turgor pressure is a fast (propagation equal to the velocity of sound in the fluid – soap – medium) and unperturbed (isotropic) messenger in the time-evolving cell. Moreover, hypotonic or hypertonic treatment of the periodically elongating cells of pollen tubes led to the recognition that the growth (growth rate) oscillation frequency was “doubled” or “halved” with respect to the unperturbed (isotonic) mode (ibid.). The oscillation frequency in such a system has been determined (Pietruszka, 2013) to obey the following relation: ω ~ √P, which means that the oscillation growth angular frequency is directly related to the pressure.

From these brief accounts, the question arises – What drives plant cell growth – cell wall loosening or osmotic pressure? Even though there is consent about the role of cell wall properties and turgor pressure in growth, a discrepancy about what drives the initial episode of cell extension exists (Zonia and Munnik, 2007). One model proposes that cell wall loosening occurs first and permits the expansion of the cell due to decreased wall rigidity. There is considerable evidence to support this view (Cosgrove, 2005b). However, cytomechanical studies have revealed that the visco-elastic properties of the pollen tube cell wall in the apical growth zone do not change considerably (Geitmann and Parre, 2004) during growth oscillations. Hence, it was suggested that the cycles of cell wall loosening cannot be the driving force of oscillatory growth (Zonia and Munnik, 2007). Moreover, it was shown that increased pressure induces the expansion of the cell wall and drives growth in *Chara* cells (Proseus et al., 2000; Proseus and Boyer, 2006). An intermediate LOS model was put forward (Wei and Lintilhac, 2003; Wei et al., 2006) that proposed a gradual increase of turgor pressure to a critical point, P_c_, at which the loss of stability initiates wall extension and growth. Without going further into the details of the ongoing discussion (e.g. Zonia and Munnik, 2007; Winship et al., 2010; Winship et al., 2011) and bringing together these different opinions, we propose a chemical potential based scenario. In what follows, we have intentionally omitted irrelevant details in order to make the main steps clear when introducing it and to build a story with a logical flow.

### Results

#### Mystery of the Lockhart coefficient explained?

If one imagines that the growth rate, as a function of pH, has a (Euler) beta function form for any fixed temperature (*T*) and a similar dependence as a function of *T* for a fixed pH (see SI Table 1 and SI Figures 1 and 1.1), one can conclude that the volumetric growth rate is proportional to the product of the beta functions of *T* and pH (and to the ‘effective’ turgor pressure *P – Y*). This assertion can be the starting point for the ‘derivation’ (in the first approximation in which we neglect the nested pH(μ_H+_(T)) dependence) of the equation below. Simply speaking, the above statement for the relative growth rate can be expressed in a way that is analogous to the Lockhart (1965) equation as an Ansatz (“educated guess”) and can be postulated in the general form

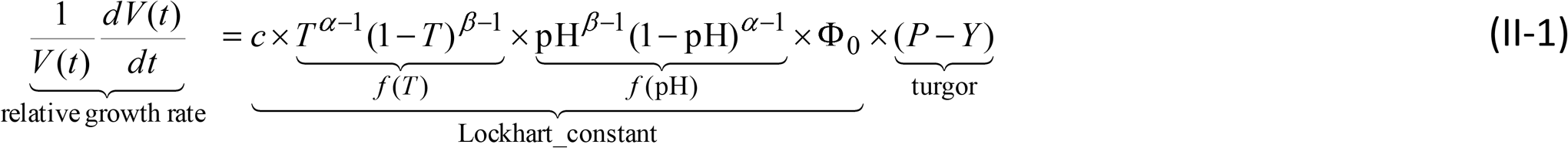

at a constant turgor pressure *P* and yield threshold *Y* (both in [MPa]), where c is a scaling constant. We assume concomitant water uptake; note that the *P – Y* term can be substituted with a more appropriate turgor plus osmotic pressure term in order to allow for water transport (aquaporins) due to the changes in the osmotic and water potential (Obermayer, 2017 in *Pollen Tip Growth*)). *V* = *V*(*t*) is the increasing volume of a growing cell over the course of time. Here, “pH” is treated, as before, as a non-separable variable name and a constant Ф_0_ was introduced for dimensionality reasons – [Ф_0_] = MPa^−1^ × s^−1^, as in the Lockhart equation. In isothermal conditions, the integration of Eq. (II-1) yields the solution for the enlarging volume in the form

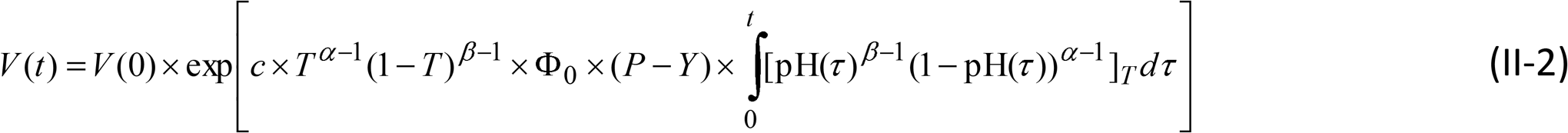

where 0 and *t* denote the initial and final time, and c is proportional to the reciprocate of the product of the beta function normalisation constants (see Eq. (I-1); it can be incorporated to Ф_0_). Note that, unlike the Lockhart equation, the “acidification” that is present in Eq. (II-2), which is represented by the value of pH, prevents a plant cell from unlimited exponential growth (burst). In contrary, the solution of the Lockhart equation *V*(*t*) = *V*(0)exp(Ф_0_(*P – Y*))*t* means that plants grow with no limitation, which is an apparent contradiction to real life phenomena. Equation (II-2) correctly delivers not only the initial phase of slow, and next of rapid growth, but also the deceleration and saturation phases (Fig. 6). Equation (II-2) holds for dynamic pH (in Fig. 7 changes in the volume and saturation effect depend on the simulation parameters).

**Figure 6.**
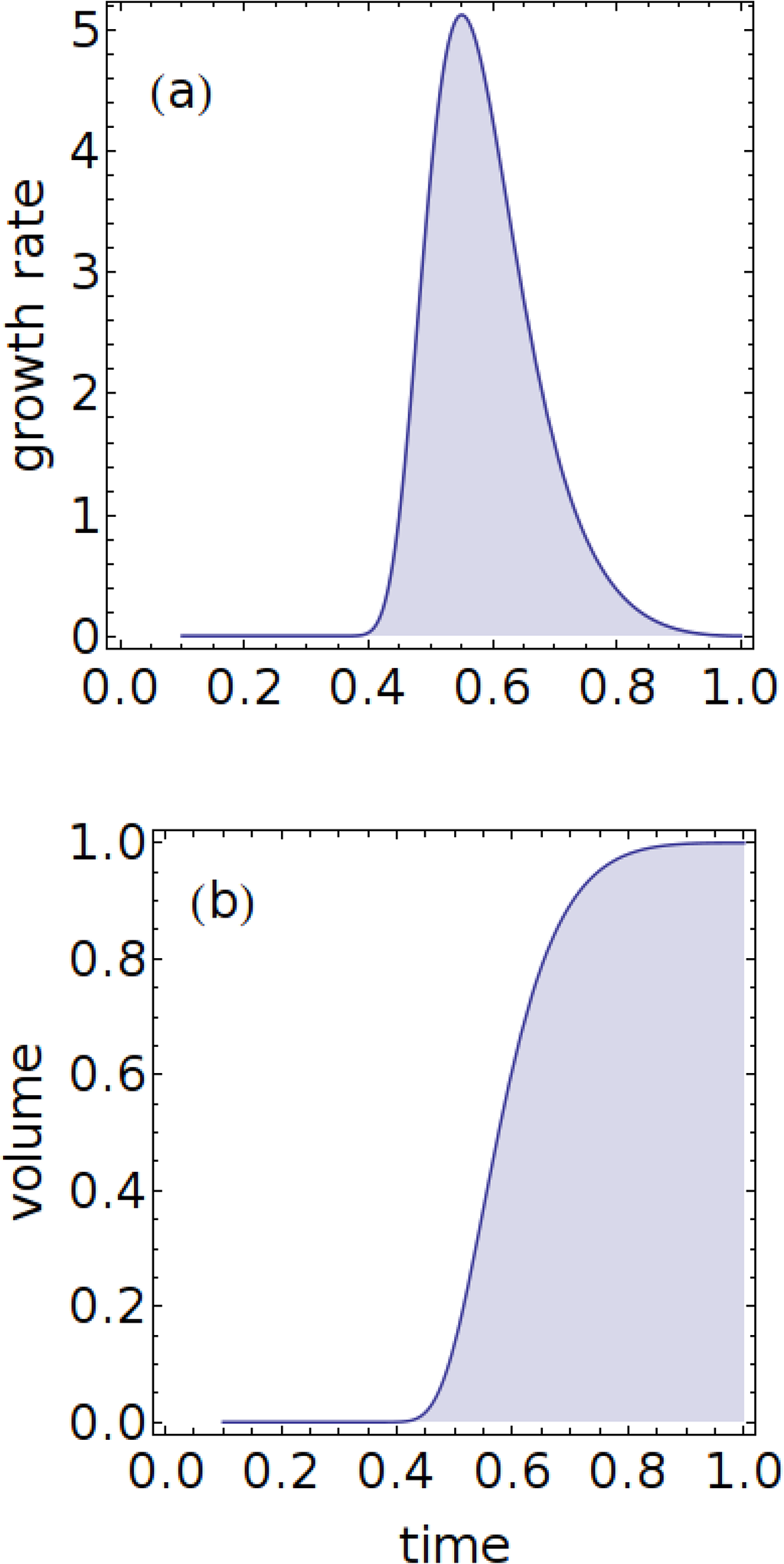
(a)Growth rate plot obtained for Eq. (II-1) where pH was assumed to be inversely proportional to time (Lüthen et al. (1990), Fig. 6). Simulation data: α = 3, β = 2; arbitrary units. (**b)** Typical growth curve obtained from Eq. (II-2) – all growth phases are clearly visible: initial phase of slow growth, accelerated growth, quasi-linear phase, deceleration and saturation (Fogg, 1975). See also Fig. 7 for parameterisation of Eq. (II-2) with respect to pH (7a), temperature (7b) and turgor pressure (7c).

By recalling the canonical form of the Lockhart equation with the extensibility Ф controlling the rate of growth (proportional to the excess of turgor pressure *P – Y*)

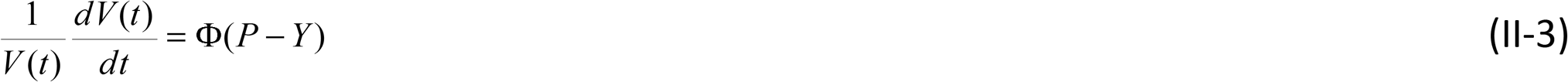

and comparing Eqs. (II-1) and (II-3), we obtain the estimate for the Lockhart ‘constant’:

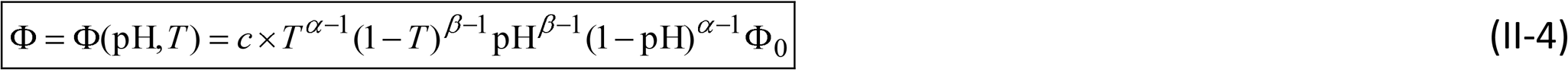

where the exponents α and β are numbers that are independent of pH (or μ) and *T*, whereas pH = pH(*t*). Note that Eq. (II-1) turns into its canonical – Lockhart – form, Eq. (II-3), for α = β = 1 and c = 1. Hence, the Lockhart coefficient Ф, which is responsible for visco-plastic extensibility, is an analytic function of pH and T on the physically accessible interval from 0 to 1 for each variable (Part I). However, the augmented form of the Lockhart equation, which is expressed by Eq. (II-1), provides the possibility to not only describe monotonic cell expansion correctly (Figs 6–7), but also the oscillatory mode of the extension of the pollen tubes. Apparently, in the case of oscillatory turgor pressure, the additional (visco-elastic) term 1/ε × d*P*/d*t* (Ortega, 1985; Sridhar et al., 2018) can be introduced (added) into equation (II-1) on the right side for completeness. However, another proposal may include a variability of *P* in agreement with pH changes in a self-consistent solution of Eq. (II-1). This, in turn, can lead to the growth rate oscillations that are observed in pollen tubes. Additionally, the remarks concerning the possible changeability of *Y* during growth (Pietruszka, 2012; Sridhar et al, 2018) are relevant in the higher order of approximation.

**Figure 7.**
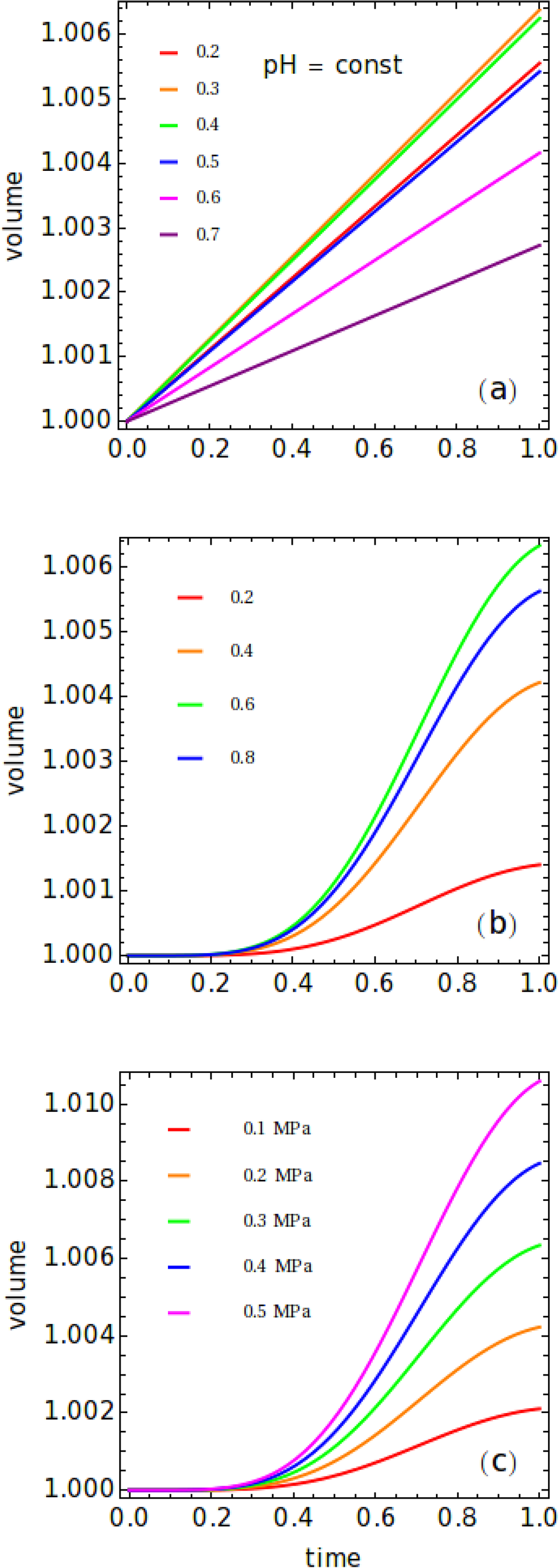
Volumetric growth calculated from Eq. (II-2), an exemplary fit. **(a)** At constant pH. *P – Y* = 0.3 MPa, *T* = 0.6 (~ 25ºC); pH-scaled values (0.2 – 0.7, increment 0.1) are shown in the plot (*T* and pH scaled to [0, 1] interval for all charts – see text). Note the linear character of the solution of Eq. (II-2) for pH = const and the maximum expansion for the normalised value of pH ~ 0.3 – 0.4 (note that the slope of the volumetric extension does *not* increase sequentially with decreasing pH). **(b)** For changing temperature *T*. Simulation parameters: P – Y = 0.3 MPa, T = 0.2, 0.4, 0.6, 0.8 (legend); pH decreases like ~ 1/*t*. Note the typical sigmoid-like curve solution of Eq. (II-2) over a [0, 1] time interval that corresponds to a rapid (~ 1/*t*) pH drop. Maximum extension (rate) is observed approximately for *T* = 0.6 (~ 25 ºC) and decreases at both higher and lower temperatures as is expected. **(c)** For changing effective turgor pressure *P – Y*. Simulation parameters: *P – Y* = 0.1, 0.2, 0.3, 0.4, 0.5 MPa (legend); *T* = 0.6 (~ 25ºC). Note the sigmoid-like growth curve for a rapidly decreasing pH. The growth curves increase with pressure in a sequential order. The simulation parameters that are common for all of the figures: α = 3, β = 2.

Note that Eq. (II-1) guarantee that the maximum growth rate is for the maximum of beta *f*(α, β, *T*) function. It is also evident that oscillations of pH ≈ sin(Ω∙*t*) will produce oscillations of the exponent function, however shifted in phase like cos(Ω∙*t*). This, in turn, can be responsible for – as yet unexplained – about 90 degrees phase shift observed by many authors in pollen tubes (e.g. Messerli et al., 1999).

Calculation of the α and β shape coefficients for the actual empirical data is commented below.

#### Determination of the α and β shape coefficients for experimental data

Potential applications of our formalism lie in the α and β parameters and the direct use of pH(t) data in calculating the volume increase d*V*. However, in Eq. (II-2), we have exp(*f*(α, β, c) × integral (α, β)) – such problems have several local extremes. Therefore, the solution of Eq. (II-1) belongs to the class of highly nonlinear optimisation problems and will be presented in a separate paper.

We suppose that the biological meaning of the α and β shape exponents that is not discussed here may have meaning for more abundant sets of data. Nonetheless, it seems that Eq. (II-2) can be treated as the transformation 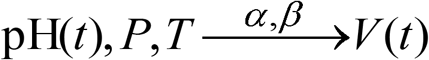 for the increasing volume of a plant cell (corresponding to the pH-driven cell enlargement).

#### Monotonic cell elongation

In order to check our Ansatz, which is expressed by Eq. (II-1), we also examined the decreases of pH that were induced by auxin (IAA) and fusicoccin (FC) in Lüthen et al. (1990), which is presented in Fig. 6 therein. There, the additions of IAA and FC resulted in rapid drops of pH. The authors asserted that these lower pH values are responsible for the rapid increase in the growth rate. We digitised the pH data that was presented by Lüthen et al. (1990) and inserted these (point by point) into Eq. (II-2) separately. As a result, we received two fragments of growth curves, which are presented in Fig. 8. (Note that to determine the cell volume evolution over time, we only needed the values of the following dynamic variables: pH, temperature and turgor pressure; the initial volume V_0_ was needed for scaling; T = const. and P ≈ const. though this may also change in time). Interestingly, the simulation data for the α and β parameters, which – represent the properties of the wall, were adapted from Part I by combining the EoS with the evolution equation (II-2). These data, for which Eq. (II-2) is *not* divergent (unlike the Lockhart equation), closed the “system of equations” for growth. By the “system”, we mean (a) the EoS, which describes the properties of the cell wall irrespective of the extension mechanism and (b) the equation of time evolution, Eq. (II-2), which describes the dynamics of the growing system at temperature *T* (see also SI Figure 6 and Pietruszka, 2012). Moreover, the direct use of Eq. (II-2) to our own experimental data is presented in Fig. 9, where pH(*t*), *P* and *T* transform directly into the elongating height (volume *V*) of a plant tissue (note the correctly calculated initial value of 1).

**Figure 8.**
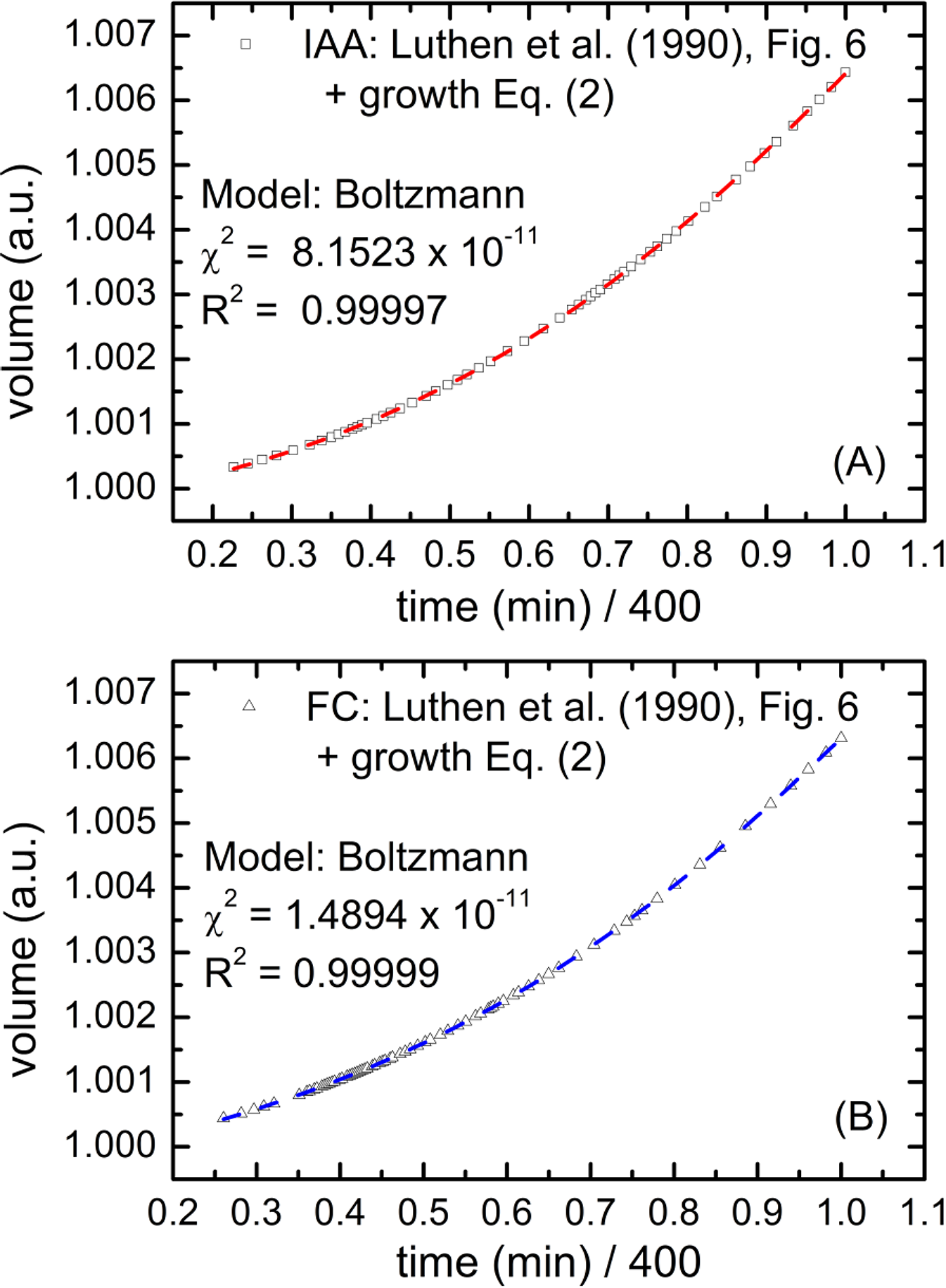
Effect of an external decrease of pH on the accelerated phase of the expanding volume of maize coleoptiles as obtained by the pH drops that were induced by 10^−5^ M IAA (A) and 10^−6^ M FC (B) in an experiment conducted by Lüthen et al. (1990). Squares and triangles – converted (recalculated) separately (point by point) using Eq. (II-2) from the digitised data presented in Fig. 6 by Lüthen et al. (1990); dashed curves – sigmoid curve (Boltzmann) fit (Microcal Origin). Conversion data: α = 3 and β = 2. Simulation parameters: *T* = 0.62 (~ 26 oC) and *P – Y* = 0.3 MPa.

In this paragraph, we demonstrated that equation (II-2) correctly describes monotonic growth in the context of the “acid growth hypothesis” (note that pH is explicitly included in this equation; see also Figs 8 and 9 and Eq. (9) in Pietruszka and Haduch-Sendecka (2016) and the conclusions therein), which validates its applicability not only for fusicoccin, but also for auxin (Hager, 2003). The high accuracy of the theoretical fits (solid lines) with those that originated experimentally (Fig. 6 in Lüthen et al., 1990) or especially to our own data, which is presented in Figs. 8 and 9, respectively, strongly support the presented approach. However, the question about the possible applicability of Eq. (II-2) to oscillatory growth still remains.

**Figure 9.**
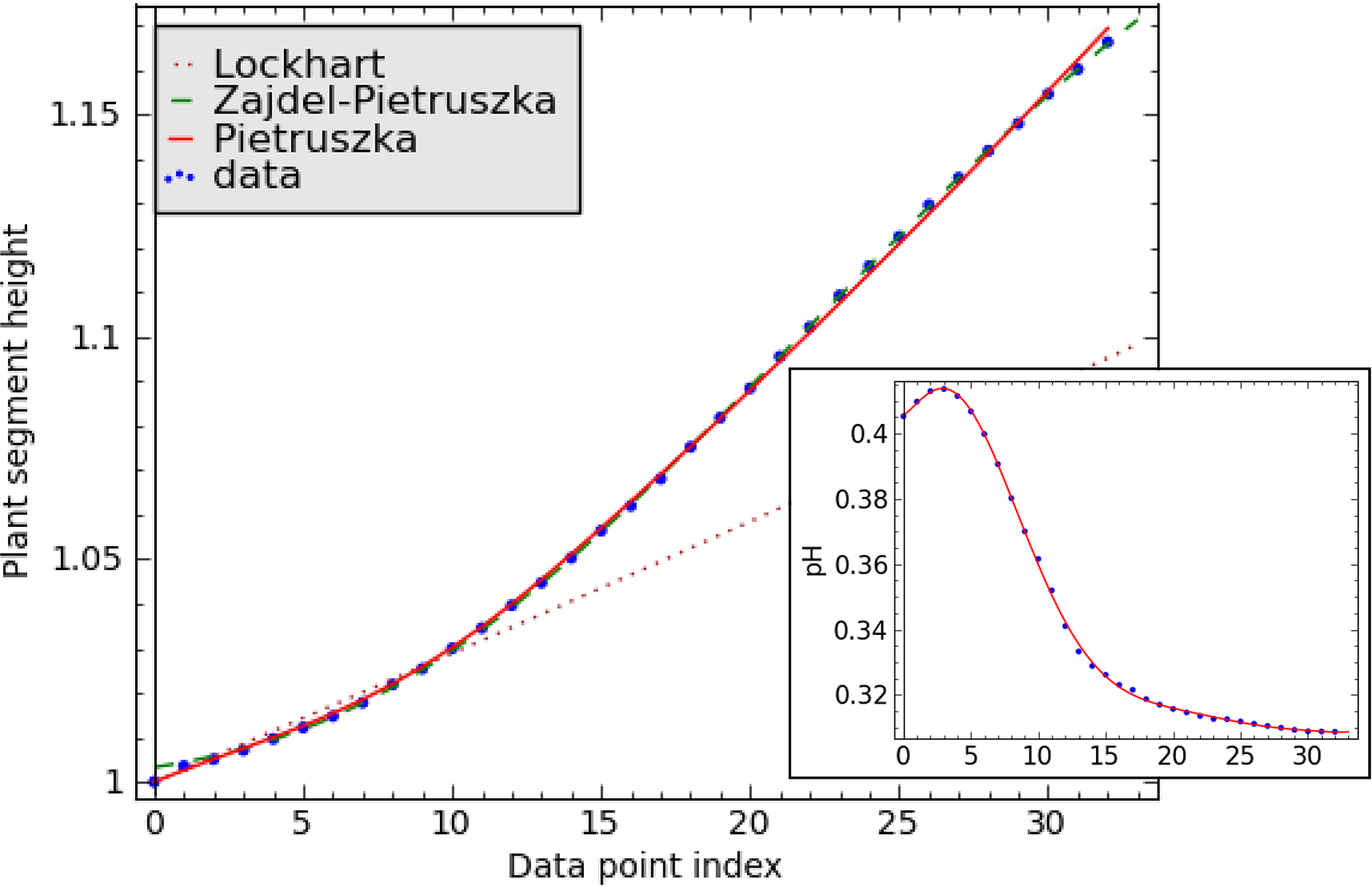
Effect of a decrease in pH on the elongation growth [cm] of a maize coleoptile segment (inset: blue dots – discrete data from an APW-driven experiment, red line – interpolation by a polynomial of the 5^th^ order). Orange dotted line – Lockhart (1965); green dashed line – experimental values interpolated by Zajdel-Pietruszka double-exponent (Zajdel et al., 2016); red line – calculated from Eq. (II-1), in collaboration with Jerzy Kosek. Simulation parameters: *T* = 0.6 (~ 25ºC) and *P – Y* = 0.3 MPa; data point index corresponding to 15 min = 900 s intervals. An example of the direct use of the pH(*t*), 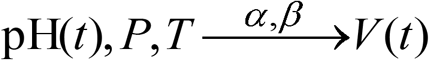 conversion, Eq. (II-2), for the increasing volume of a plant cell (provided the segment cross-section area is included).

#### Periodic cell elongation

It is known that the group of oscillating variables during growth includes at least ion fluxes (Sukhov et al., 2011) and internal free concentrations of Ca^2+^, Cl^−^, H^+^ and K^+^ ions (Holdaway-Clarke et al., 1997; Holdaway-Clarke and Hepler, 2003), the cytoskeleton (Geitmann, 2010), membrane trafficking (Feijo, 1999; Feijo et al., 2001; Zonia and Munnik, 2008) and cell wall synthesis (see Portes et al., 2015 and Hemelryck et al., 2017 for review). However, despite the progress made in this domain, the central core-controlling mechanism is missing (Damineli et al., 2015). Moreover, how it produces a macroscopic outcome in the form of apical growth also remains unknown. A possible link between this oscillatory component at the molecular level and the structurally (spatial arrangement) and temporally organised apical (periodic) growth is proposed below.

The pH value is a logarithmic measure of the H^+^ activity (the tendency of a solution to take H^+^) in aqueous solutions. However, pH can be directly expressed in terms of the chemical potential (μ_H+_) of protons through the relation

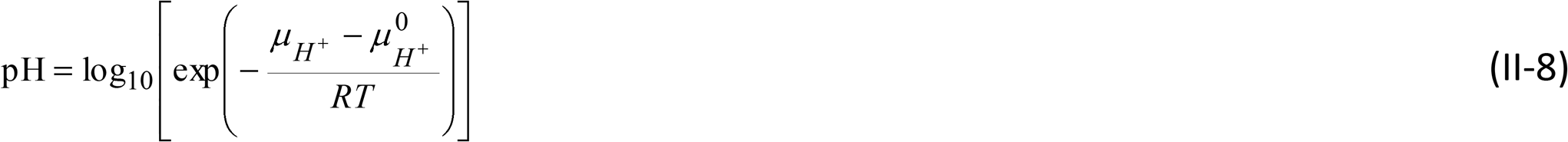

where R denotes the gas constant, *T* stands for the absolute temperature and 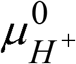 is the reference potential. In this context, pH is not a typical generalised coordinate, although the microscopic state of the system can be expressed through it in a collective manner. Just as temperature determines the flow of energy, the chemical potential determines the diffusion of particles (Baierlein, 2001).

After inserting Eq. (II-8) into Eq. (II-2), we can calculate the influence of the chemical potential oscillations of H^+^ ions, μ_H+_ = μ_H+_(*t*), on the volumetric expansion. A simplistic numerical simulation reveals that the volumetric expansion represents a typical growth (sigmoid) curve (like the one presented in Fig. 6b). However, in this case, when superimposed onto the simulated oscillations of the chemical potential, they produce the growth rate oscillations in a straightforward manner (SI Figure 4 and Pietruszka and Haduch-Sendecka, 2015a). This type of oscillations was observed in the rapid expansion of pollen tubes (Plyushch et al., 1995; Hepler et al., 2001; Feijo et al., 2001), and the source of this periodic motion has been the subject of a long-lasting debate among plant biologists. Portes et al. (2015) noted that “Since the ultradian rhythms present in pollen tubes are not temporally coincident with protein synthesis and degradation, a simpler mechanism might be triggering the overall oscillation”. Further, the authors claimed that “the strongest candidates seem to be the membrane trafficking machinery and ion dynamics, since they are the only candidate mechanisms that may oscillate independently of tube extension (Parton et al. 2003)”. Indeed, in what follows a chemical potential-induced (ultimately connected with ion dynamics since it measures the tendency of particles to diffuse) recurrent model, which triggers the overall oscillation, is proposed to explain this oscillatory mode of a single cell extension.

### Discussion

Indirect measurements of the chemical potential (μ_H+_) using pH can be compared to our previous measurements of the electromotive force (EMF) in order to detect phase transitions, which are localised by the ‘kinks’ in the chemical potential in condensed matter physics (Matlak and Pietruszka, 2001; Matlak et al. 2001). Such ‘kinks’ in the chemical potential always localised the occurrences of different types of phase transitions (where the change of state of a macroscopic system occurs) such as ferromagnetic, anti-ferromagnetic, superconducting, re-entrant or structural phase transitions (Matlak et al., 2000). Based on these observations and considerations on the EoS for plants, we can combine these outcomes together in a single coherent account.

#### Chemical potential-induced ratchet-like growth in plants

Launching our reasoning from the EoS and putting the differing approaches that were outlined in the Introduction together, we can construct a plant cell expansion recurring model. The latter, though simplistic for obvious reasons, not only describes one cell growth cycle in physical (mechanistic) terms, but also demonstrates how subsequent growth cycles can arise and even how the entire process may eventually decay (compare with Rounds et al., 2010).

The hypothetical steps in the transport of cations (such as H^+^) against its chemical gradient by an electrogenic pump and its consequences are presented in Figure 10 and SI Figure 5. These can be described as follows:

A. The initial state of a growing cell (we assume the chemical potential μ = μ_0_ for reference only).
B. The outward, active proton transport across the plasma membrane interface creates gradients of pH or equivalently gradients of the chemical potential μ = μ_H+_, which drives the transport of ions and uncharged solutes. The plasma membrane (marked by a vertical line) depicts the boundary between the cell wall and the cytoplasm.
C. H^+^-ATPase activity regulates cytoplasmatic pH (and controls cell turgor pressure). At the microscopic level, we can observe a singular behaviour at a certain critical value of the chemical potential μ = μ_c_ (see also Fig. 5c). This chemical potential activity can be interpreted as a signature for an ST that is occurring in the cell wall – during an ST of a given medium, certain properties of the medium change, often discontinuously, as a result of a change in some condition, here: μ.
D. The increased involvement of H^+^ ions in cell wall building and exocytosis (Holdaway-Clarke and Hepler, 2003) by δ*V* raises the turgor pressure inside the cell by δ*P*, thus producing force *F*, which may cause cell wall loosening and elongation by δ*l* (see Wei and Lintilhac, 2007 for a loss of stability approach and Sridhar et al., 2018 for a statistical model of cell wall dynamics). The physical quantity that may be responsible for signal transduction here is pressure (e.g. Zonia, 2010), which is a scalar quantity.
E. The increased pressure at the apical plasma membrane by δ*P* induces the stress relaxation of the cell wall, relaxation-driven cell expansion by δ*V* and triggers the start of the next growth cycle with a slightly modified turgor pressure ~ *P* – δ*P* (see the linearly descending pressure plotted in Fig. 1 in Benkert et al., 1997). The system restores the initial configuration and allows a new growth cycle to begin (explicit in pollen tube oscillations). The latter means it is reset at the end of the cycle (Hemelryck et al., 2018).

**Figure 10.**
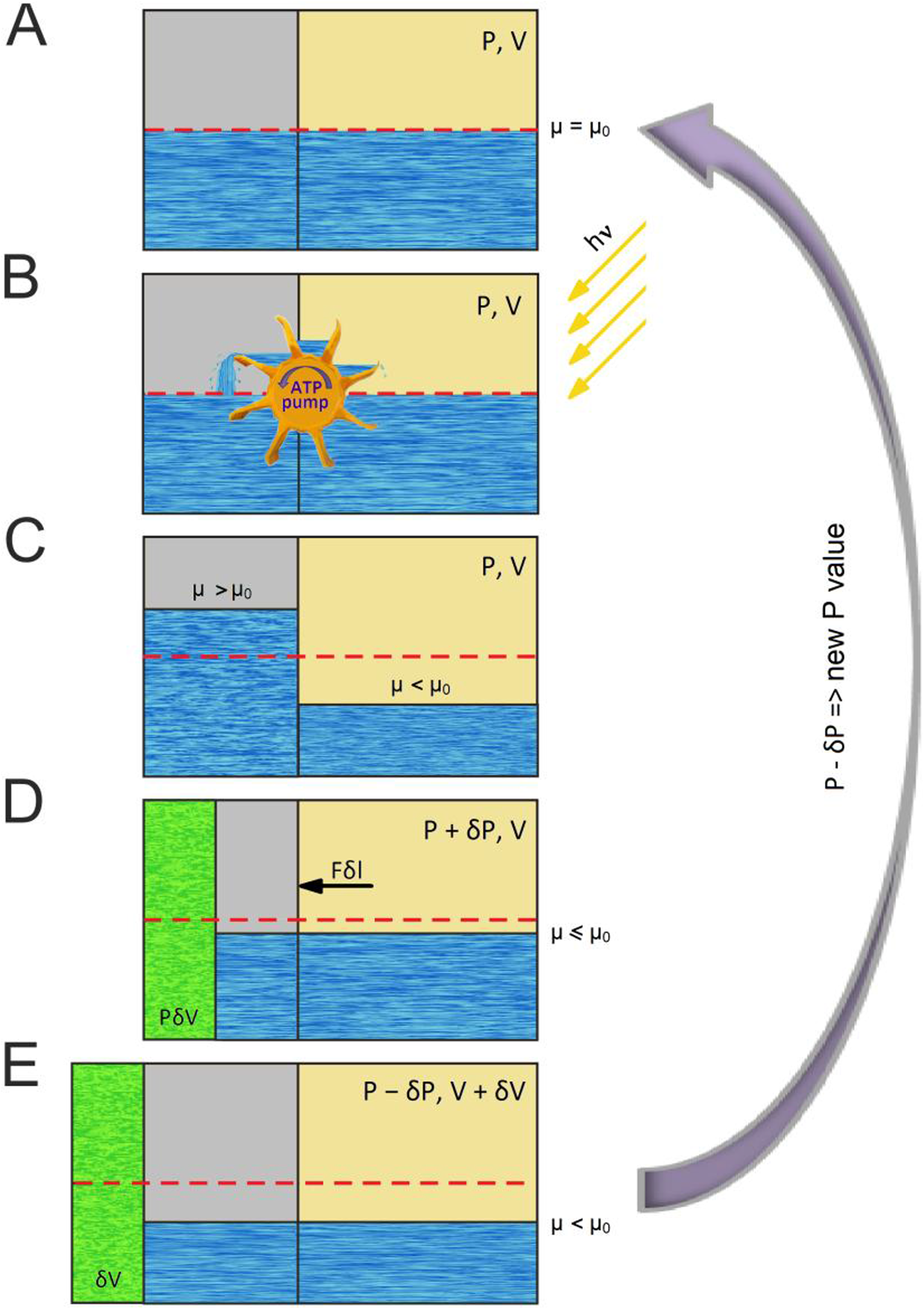
Plant cell expansion recurring model – schematic view (though, in general, it should be solved self-consistently). **A.** Initial state: the chemical potential μ (μ_0_ – reference potential) equilibrated in both compartments; turgor pressure equals P; cell volume equals V. **B.** Energy (E = hν) consuming proton pump driven by the hydrolysis of ATP transport protons (ions) against the electrical gradient. **C.** Chemical potential difference is established between the compartments – the system is far from equilibrium; turgor pressure equals *P*; cell volume *V*. **D.** Synthesis of cellulose at the plasma membrane and of the pectin and hemicelluloses components with a Golgi apparatus depositing a layer on the inside of the existing cell wall, thus causing an increase of the cytoplasm pressure by the infinitesimal value of δ*P*: *P* + δ*P*(*t*), though the cell volume still remains *V*. **D.** New wall synthesis. Elevated turgor pressure *P* by a δ*P*(*t*) amount (Benkert et al., 1997) produces force *F*, which acts on the wall (energy stored *P*δ*V*). The chemical potential starts to drop due to the overall particle loss in the system. **E.** Increasing turgor pressure releases energy to extend the existing cell wall chamber by δ*l*; the chemical potential further decreases before the next cycle (arrow). After the completed growth cycle, turgor pressure diminishes by the δ*P*(*t*) value due to leakage or the other mechanisms illustrated in the text. Another growth cycle begins (arrow), however, with a new initial value of *P* equal to *P* – δ*P*(*t*), *V* equal to *V* + δ*V* and μ equal to μ – δμ (exaggerated). **Legend.** Left compartment: plasma membrane (PM) and cell wall. Right compartment: cytoplasm. Grey – PM, blue – H^+^-”Fermi sea”, green – new cell wall.

As a result, the problem of causality, which was mentioned in the Introduction, is resolved in a <u>self-consistent manner</u>, thus potentially leading to a paradigm change that can be expressed by a *self-consistently evolving recurring model*. Obviously, the latter should be further examined by numerical calculations (self-consistent solutions at subsequent time intervals) and confronted with the existing representations (Baskin, 2015, see Fig. 1.1 therein for a complementary Maxwell-type model). The possible mechanisms of the participation of pH changes on photosynthesis (Grams et al., 2009; Sukhov et al., 2014; Sherstneva et al., 2015; Sukhov et al., 2016) can also be included in the form of a phase-locked loop, thus closing the system of equations. The existence of oscillating behaviours in the investigated system, however, implies the existence of a control by feed forward or feedback loops, which are inserted into a self-regulating mechanism (Portes et al., 2015). The latter statement can be translated in mathematical terms into a self-consistent evolutions equation that corresponds with growth.

We will briefly discuss the major novelty of the recurrent model as compared to the earlier work in the context of the previous models. The three main approaches that were outlined in the Introduction to Part II emphasised a single attribute that is responsible for expansion (or periodic) growth. However, it seems that none of these factors alone (such as oscillatory wall synthesis, excessive turgor pressure or turgor pressure oscillations) can explain all of the observable modes of cell expansion and growth. Moreover, because of the intricate dependence between the processes that are involved in tip growth, causal relations cannot be easily inferred solely from phase relations (Pietruszka and Haduch-Sendecka, 2015b) as simultaneous information about the role of the participating ions and the actual growth rate amplitude are required (ibid).

Furthermore, intracellular ion dynamics has been shown to play an essential role in pollen tube growth (Portes et al., 2015), while calcium, protons and chloride are also considered to be of major importance. Because the [Ca^2+^]_i_ pool at the apex oscillates (ibid.), it has also been considered as supporting growth (Messerli and Robinson, 1997). Additionally, it has been experimentally verified that pollen tubes stop growing when the [Ca^2+^]_i_ gradient is dissipated (Pierson et al., 1994). An interesting fact regarding chloride dynamics is that in contrast to Ca^2+^ and H^+^, the ionic fluxes occur in the reverse direction at the apex, thereby elevating the chemical potential gradients even further.

Protons have received much attention as they play a crucial role in pollen tube growth. Since water, which ionises spontaneously, is the major component of living cells, H^+^ ions are known to be involved in enzymatic activity, endo/exocytosis and many other processes (Portes et al., 2015 for review). Therefore, pH inside cells has to be regulated in harmony with the growth phases (Messerli and Robinson, 1998). It was found that pollen tubes showed an acidic domain that was located close to the membrane apex – the latter was present only during growth. In addition, the localisation of pollen-specific H^+^-ATPase provides evidence that indicates correlations with the proton extracellular fluxes and the acidic domain at the apex (Certal et al., 2008). It looks as though proton dynamics is involved in the mechanisms for maintaining the polarity that ensures pollen tube growth (Portes et al., 2015). Moreover, it has been established (Agudelo et al., 2016) that the pollen tube response is correlated with the conductivity of the growth medium under different AC frequencies, which is consistent with the notion that the effect of the field on pollen tube growth may be mediated *via* its effect on the motion of ions. This is, however, in accord with the presented chemical potential (ratchet) scenario and the representative time series of the oscillatory traces of incoming cations (H^+^ and Ca^2+^) or outgoing anions (Cl^−^) at the growing tip. Hence, from the point of view of the recurrent model, the equilibration of the chemical potential for all of these species should also be the major mechanism for the cessation of growth.

Ion homeostasis and signalling are crucial in regulating pollen tube growth and morphogenesis and affect upstream membrane transporters and downstream targets (Michard et al., 2016). Pollen tube growth is strictly dependent on ion dynamics. Ion fluxes and cytosolic gradients of concentration have been mechanistically associated with the action of specific transporters, especially protons (ibid.). In the recurrent model put forward in this section, though extremely simplistic, H^+^ chemical potential dynamics may serve as a “still missing” (Portes et al., 2015) central core-controlling mechanism (pacemaker) that is able to produce a macroscopic outcome, i.e. structurally and temporally organised apical growth, especially since H^+^ usually oscillates with the shortest (basic) period.

On the other hand, the short period oscillations of pH, which induce changes in wall properties, can be translated into the volume pulsatile extension by Eq. (II-1). Provided that cross-correlation (vs. time) of pH and P exists, turgor pressure P should vary slightly (Fig. 1 in Benkert et al., 1997) in accord with minute pH drops (note the pressure linear decrease, ibid.); equation (II-1) allows for such analysis. This should also explain the observed phase delays in growth rate and ionic fluxes in pollen tubes in mechanistic terms (e.g. Messerli at al., 1998). This is, however, beyond the scope of this work and should be analysed in another study.

### Summary

This aim of section was to develop a unifying model to simulate plant cell expansion and growth. In the chemical potential (μ) representation, growth can be treated as a dynamic series of STs that can take place in the primary plant cell walls (Part I) during subsequent time intervals. We suppose that such changes would proceed, e.g. from an ordered to disordered state, from a stressed to relaxed state, from a “covalent bonds state” to “disrupted covalent bonds state” or even through more subtle pathways in the wall polymer network, which themselves constitute an intriguing research task. Furthermore, the singular causality, whereby only biochemical wall loosening can lead to stress relaxation and turgor loss, is replaced by a self-consistent recurring model, in which all of the prevailing variables constitute the macroscopic evolution of the plant cell. A common denominator appears to emerge that ties all of the growth factors together, namely the chemical potential.

At the very beginning of this section we asked, whether the volumetric growth of plants could be calculated from pH, T and P. It turned out that the answer is “yes”, provided that it is delivered *a posteriori*, i.e. the pH growth data is already known. This reasoning, however, can be reversed – by knowing pH(t), T and P, we can predict the cell volume V expansion over time using model equation II-2.

As we have learned, from the physical point of view, cell volume expansion in plants follows the *V*(*t*) = *f*(*P*, *T*, pH(*t*); α, β) analytic transformation through the coupling between the mechanical (pressure), thermal (temperature) and chemical (pH, μ) degrees of freedom. In this process, the chemical potential of the intervening ions can play an important role – distant (nonlocal) correlations are no longer controlled externally, but by the system itself, which is similar to the self-organised critical states (Bak et al., 1988, Hesse and Gross, 2014) and remains in agreement with the statistical approach (viscoplastic and viscoelastic terms mapped onto a statistics of independently acting elastic springs) of Sridhar et al, 2018. Indeed, it has been suggested that a growing cell proceeds along a critical trajectory over 1/*f* edge (*f* – fluctuation frequency of ionic channels), which is typical for regular growth (Pietruszka, 2018). The latter observation can be confirmed by the cross-correlations that are maximum for the control (normal) growth, see Figs 4–5 in Olszewska et al., 2018. Because of the pH(μ) dependence, the signalling cascade during the growth of a cell seems to be triggered and controlled by the chemical potential.

### Conclusions

By resolving the duality of low pH or auxin action (producing acidic pH) against temperature, not only have we introduced an EoS for the realm of plants, but also, by considering wall extension growth as a dynamic cascade of chemical potential driven STs, we have identified the critical exponents for this phenomenon, which exhibit a singular behaviour at a critical temperature and critical pH in the form of power laws. A common unifying principle that can be applied for either condensed matter (studied in physics) or living matter (a subject of biology) is proposed for all of the chemical potential driven transitions.

## Author Contributions

MP conceived the ideas, conducted the analyses, conducted the calculations and estimates and wrote the manuscript.

